# Validation of cross progeny variance genomic prediction using simulations and experimental data in winter elite bread wheat

**DOI:** 10.1101/2023.09.26.558758

**Authors:** Claire Oget-Ebrad, Emmanuel Heumez, Laure Duchalais, Ellen Goudemand-Dugué, François-Xavier Oury, Jean-Michel Elsen, Sophie Bouchet

## Abstract

The utilization of genomic prediction is increasing in crop breeding to parental selection and mating. The Usefulness Criterion (UC) that considers Parental Mean (PM), progeny Standard Deviation (SD) and selection intensity has been shown to increase the likelihood to get outstanding progenies compared to mating using PM alone while maintaining more diversity in the germplasm for next generations.

This study estimates our ability to predict UC and its two components (PM and SD) using simulations and experimental data (73-101 winter bread wheat crosses depending on the trait, with 54.8 progenies on average) including heading date, plant height, grain protein content and yield evaluation. The training population comprises 2,146 French varieties registered during the last 20 years and INRAE-AO breeding lines.

According to simulations, prediction ability increases with heritability and progeny size and decreases with QTL number, most notably for SD. We used as a reference a TRUE scenario, *i.e*. an optimal situation where TP is infinite and where marker effects are perfectly estimated. SD was strongly impacted by the quality of marker effect estimates. In simulations, considering the error in marker effect estimates improved SD predictions for quantitative traits with low heritability. In experimental data, the interest of this method was limited.

PM and UC were reasonably predicted for all traits, while SD was more challenging. This pioneering study experimentally validates genomic prediction of progeny variance. The ability of prediction depends on trait architecture while the realization of cross potential in the field necessitates a sufficient number of progenies.

**Key message:** From simulations and experimental data, the quality of cross progeny variance genomic predictions may be high, but depends on trait architecture and necessitates sufficient number of progenies.

## Introduction

How humans choose candidates for selection has evolved over time. At first, selection based solely on phenotypic observation of progeny, was next assisted by pedigree-based predictions, more particularly in animals (Henderson 1975; Falconer and Mackay 1996). Nowadays, genomic predictions of individual genetic values (Whittaker et al. 2000; Meuwissen et al. 2001) and of cross values (Schnell and Utz 1975; Zhong and Jannink 2007; Lehermeier et al. 2017; Danguy des Déserts et al. 2023) are promising and open up new prospects for improving crossbreeding schemes and optimizing long-term genetic gain (Allier et al. 2019a, 2019b).

In plants, crosses are essentially selected to secure a high mean yield performance of the progeny. Breeders seek to identify which crosses will produce the best superior progeny in order to ensure genetic gain in the next selection cycle, and to maintain long-term selection gain. It is thus important that the selected crosses also generate a high genetic progeny variance. A strategy to find these crosses is to select on the Usefulness Criterion (*UC*) (Schnell and Utz 1975; Zhong and Jannink 2007) defined as: *UC* = *PM* + *i* × *SD*, where *PM* is the Parental Mean, *i* the selection intensity corresponding to the fraction *q* of selected progenies, and *SD* the progeny Standard Deviation (square root of progeny variance). PM can be estimated from the mean of additive parental genetic values using phenotypic or genomic selection. Several attempts have been made to predict progeny variance using phenotypic distances (Utz et al. 2001) and genetic distances (Bohn et al. 1999; Hung et al. 2012), but without success. According to the multiplication of recent methodological studies, genomic predictions appear to be a promising tool for optimizing complementarity between parents and breeding schemes. An algebraic formula to predict inbred progeny variance derived from a cross between two inbred lines was provided by Lehermeier et al. 2017 based on marker effect estimates and their co-segregation in progeny derived from a genetic map.

Several studies showed that selecting on criteria based on progeny variance such as UC could actually increase the genetic gain in plants and in animals (Tiede et al. 2015; Santos et al. 2019; Allier et al. 2019a, 2019b; Bijma et al. 2020). But, to our knowledge, the ability to predict the value of a cross has only been validated by simulations, no publication has yet validated the ability to predict cross values on real data.

This study was carried out to (i) estimate the ability of genomic prediction of three cross value components (PM, SD and UC) in real experimental data, and (ii) to identify the factors influencing this prediction ability based on simulations. We based our predictions on different genomic additive models (Lorenz et al. 2011) trained on a historical training population (TP) of 2,146 varieties and breeding lines of winter bread wheat. We compared these predictions to observed phenotypes of unselected progenies of 73 or 101 crosses depending on the trait.

## Materials and methods

The general idea of this study is to compare genomic predicted values of gametic variance to observed variance of progeny’s phenotypes. To do so, we first developed a simulation study by crossing real genotypes of a historical TP to evaluate the different parameters impacting the prediction ability. Then, we analyzed 4 traits of interest for winter bread wheat (yield, grain protein content, plant height and heading date) in experimental data.

### The training population

#### Datasets description

In order to predict genomic cross values, we used a TP composed of two datasets of lines from two institutes: *Institut National de la Recherche pour l’Agriculture, l’Alimentation et l’Environnement* (INRAE) with its subsidiary company *Agri-Obtentions* (AO), INRAE-AO, and *Groupe d’Étude et de contrôle des Variétés Et des Semences*, GEVES. The detailed description of the TP for the 4 analyzed traits is provided in **Supplementary Tables S1** to **S9**.

The INRAE-AO dataset was composed of a breeding population of F8-F9 winter bread wheat lines developed by INRAE-AO and evaluated between 2000 and 2022 in two French geographical area (North and South). Each year, the lines were phenotyped in up to 9 locations in France for several traits of interest including yield, grain protein content, plant height and heading date. On average, depending on the trait, between 157 and 169, and between 26 and 42 varieties were evaluated each year in North and South areas, respectively. Each variety has been evaluated for at least two years. In each location, some registered varieties were used as controls and have been evaluated for several consecutive years. We used the data base corresponding to high yield crop management (optimized pesticide, fungicide and nitrogen amount). The total number of phenotyped lines in the INRAE-AO dataset was 2,673, 2,369, 2,439 and 2,672 for yield, grain protein content, plant height and heading date, respectively.

The GEVES dataset is composed of the VATE (*Valeur Agronomique, Technologique et Environnementale* - agronomic, technological and environmental value) data on winter bread wheat collected by the GEVES as part of the evaluation of varieties for national registration between 2000 and 2022. We included in our analyses the varieties evaluated in the North and the South national bread wheat experimentation networks with high yield crop management. On average, depending on the trait, between 44 and 46, and between 26 and 27 varieties were evaluated each year in North and South networks, respectively. In these networks, some registered varieties were also used as controls and have been evaluated for several consecutive years. The total number of phenotyped lines in the GEVES dataset was 650, 644, 650 and 647 for yield, grain protein content, plant height and heading date, respectively.

#### Correlation of phenotypes between years

In order to check the reliability and consistency of observations from one year to the next in the TP, we computed Pearson’s correlations between years for each trait and each dataset (**Supplementary Tables S2** to **S9**). For the INRAE-AO dataset, the average number of varieties in common between two consecutive years ranged, depending on the trait, from 54 to 60, and from 11 to 16 in North and South areas, respectively. For the GEVES dataset, the average number of varieties in common between two consecutive years was 25 and 15 for the four phenotypes in North and South networks, respectively.

For each year, we used the following mixed model to obtain one Best Linear Unbiased Estimator (BLUE) value per line across trials:

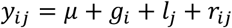

with *Y*_*ij*_ the phenotype for line *i* in location *j, μ* the grand mean, *g*_*i*_ the fixed effect of line *i, l*_*j*_ the random effect of location *j*, and *r*_*ij*_ the random effect of residual. We implemented the model using the `lme4` v1.1.27.1 R package (Bates et al. 2015). Least-squares means for each variety were then computed from this model using the `lsmeans` v2.30.0 R package (Lenth 2016). We calculated Pearson’s correlations between these values for consecutive years.

#### Phenotypes

For each line in each trial, we computed spatially adjusted means when coordinates of plots were available and arithmetic means if not. We obtained spatially adjusted means by fitting a linear mixed model where the spatial trend was modelled by means of two-dimensional P-splines using the `SpATS` v1.0.16 R package (Rodríguez-Álvarez et al. 2018). The total number of means was 57,226, 30,901, 29,445 and 44,614 for yield, grain protein content, plant height and heading date, respectively (**Supplementary Table S1**). We then calculated Best Linear Unbiased Estimator (BLUE) values for each line of the TP using the following model:

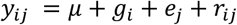

with *Y*_*ij*_ the mean for line *i* in environment (year and location intersection) *j, μ* the grand mean, *g*_*i*_ the fixed effect of line *i, e*_*j*_ the random effect of environment *j*, and *r*_*ij*_ the random effect of residual. The total number of environments was 1,031, 714, 551 and 832 for yield, grain protein content, plant height and heading date, respectively (**Supplementary Table S1**). The model was implemented using the `lme4` R package. We considered these BLUE values as phenotypes in the further analyses. The total number of BLUE values was 3,241, 2,933, 3,008 and 3,237 for yield, grain protein content, plant height and heading date, respectively (**Supplementary Table S1**).

#### Genotypes

In this study, we used 2,591 lines genotyped using the 35K Single Nucleotide Polymorphisms (SNP) representative of the TaBW280K array (Rimbert et al. 2018; Ben-Sadoun et al. 2020). After quality filtering performed with `PLINK` v1.90b5.3 toolset for minor allele frequency (filtering out variants with minor allele frequency below 0.01) and call rates (filtering out variants with missing call rates exceeding 0.4), we obtained 23,140 markers. We imputed missing genotypes with the algorithm implemented in `BEAGLE` v5.3 (Browning et al. 2018) using the genetic positions previously estimated for a West-European bread wheat population (Danguy des Déserts et al. 2021). The total number of lines with genotypes and phenotypes was 2,146, 2,062, 2,126 and 2,145 for yield, grain protein content, plant height and heading date, respectively (**Supplementary Table S1**).

#### Variance components

First, we estimated repeatability of each trait, using the following model:

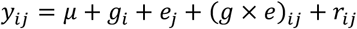

with *Y*_*ij*_ the raw observations for line *i* in environment (year and location intersection) *j, μ* the grand mean, *g*_*i*_ the effect of line *i, e*_*j*_ the effect of environment *j*, (*g* × *e*)_*ij*_ the effect of the interaction between line *i* and environment*j*, and *r* the effect of residual. All effects were considered as random. The model was implemented using the `lme4` R package. We then computed repeatability at the plot and design levels as:

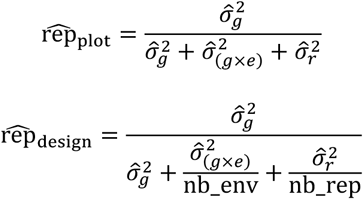

Where 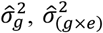 and 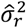 are the line, the interaction between line and environment, and the residual estimated variances, respectively, and nb_env and nb_rep are the average number of environments and replications per line, respectively.

Next, we estimated genomic heritability of each trait with a RR-BLUP (Ridge Regression Best Linear Unbiased Predictor) approach, using the following model, implemented in the `rrBLUP` v4.6.1 R package (Endelman 2011):

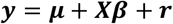

with *y* the vector of BLUE values, *μ* the grand mean, *β* a vector of random SNP effects assumed to be normally distributed 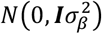, with its matrix of incidence *X*, and *r* the vector of random residual effects assumed to be normally distributed 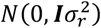. We then computed genomic heritability as:

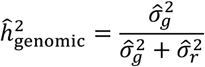

where 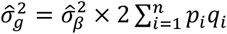, with *p*_*i*_ the allele frequency of SNP *i, q*= 1 *− p*, and *n* the total number of SNP.

#### TP Genome Estimated Breeding Values (GEBV) cross-validation procedure

We assessed the prediction ability in the TP using a 1-fold cross-validation procedure iterated 100 times. In each iteration the training set (size *N* × 0.6, where *N* is the size of the TP) is randomly sampled from the TP, and the remainder of the TP is assigned to the validation set. The prediction abilities are expressed as the average Pearson’s correlation between the GEBV and observed phenotypic values in the validation set across all iterations. This cross-validation procedure is implemented in the `PopVar` v1.3.0 R package (Mohammadi et al. 2015; Neyhart et al. 2019).

### The breeding experimental population

In this study, we generated crosses in order to phenotype their progeny so that we can observe the progeny distributions in the field and compare them to genomic predictions. Details about the experimental design are provided in **Supplementary Table S10**.

#### Data description

Within the FSOV *PrediCropt* project, a total number of 102 crosses were generated in 2020 (25 crosses) and 2021 (49 crosses) and 2022 (28 crosses). These crosses were generated by a private company (Florimond-Desprez), INRAE and INRAE-AO. The number of progenies per cross ranged from 8 to 122 (mean: 54.8 progenies per cross, **Supplementary Table S11**). The 16 crosses generated by Florimond-Desprez were evaluated in two locations (Cappelle and Houville, France), and the 86 remaining were evaluated in 4 other locations in France (Auzeville: INRAE UE GC, Clermont-Ferrand: INRAE UE PHACC, Lusignan: INRAE UE FERLUS, and Mons: INRAE UE GCIE).

In both experimental protocols (Florimond-Desprez and INRAE):

- crop management methods corresponded to high yield objectives (optimized pesticide, fungicide and nitrogen amount),
- some registered varieties and/or parents of the generated crosses were used as controls and repeated across trials (year x location),
- at least 10% of the progenies were replicated in each trial,
- each cross progeny was represented in all locations.

We phenotyped each plot of each trial for 4 traits: yield, grain protein content, plant height and heading date. Due to the unavailability of 2022 protein content data from INRAE trials, only the crosses generated in the two first years (74) were analyzed for this phenotype. The total number of plots was 1,908 and 7,570 for Florimond-Desprez and INRAE protocols respectively, representing 1,168 and 7,394 lines (**Supplementary Table S10**). We discarded from further analyses plots that were affected by lodging. After cleaning raw observations, the total number of phenotyped progenies were 5,565, 3,522, 5,591 and 5,590 for yield, grain protein content, plant height and heading date, respectively.

The progeny of each cross was examined both by genotyping a sample of offspring and by visual observation of potentially incompatible phenotypes between parents and offspring, such as wheat ear beard. Plots with incompatibilities were discarded from the further analyses.

#### Phenotypes

For each trial, we adjusted phenotypes spatial effects using the `SpATS` R package. In order to check the reliability and consistency of phenotypes between sites and years, we computed Pearson’s correlations between sites and years for each trait in each protocol (**Supplementary Tables S12 and S13**).

We then calculated BLUE values for each line using the following model:

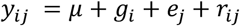

with *Y*_*ij*_ the adjusted phenotype for line *i* in environment (year and location intersection) *j, μ* the grand mean, *g*_*i*_ the fixed effect of line *i, e*_*j*_ the random effect of environment *j*, and *r*_*ij*_ the random effect of residual. The total number of environments was 11 for grain protein content and 15 for the three other traits. The model was implemented using the `lme4` R package. We considered these BLUE values as phenotypes in the further analyses. The total number of BLUE values was 5,658, 3,596, 5,684 and 5,683 for yield, grain protein content, plant height and heading date, respectively (**Supplementary Table S10**).

### Cross value components prediction ability

In this study, we considered three components for the value of a cross: the parental mean (PM), the progeny standard deviation (SD) which is the square root of gametic variance, and the UC, corresponding in our study to the expected mean of the 7% best progeny of a cross (Schnell and Utz 1975; Zhong and Jannink 2007; Lehermeier et al. 2017).

#### Algebraic genomic predictions

These three cross value components can be predicted by analytic formulae that takes into account marker effects and phase as well as recombination rates. The estimation of marker effects and the predictions of the three cross value components were performed with the `PopVar` R package. In total, we predicted the cross value components of 73 or 101 experimental crosses depending on the trait.

To estimate marker effects, we considered the following model:

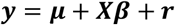

with *y* the vector of phenotypes of the TP corrected for any fixed effects (the BLUE values described previously), *μ* the grand mean, *β* a vector of random SNP effects assumed to be normally distributed 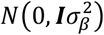, with its matrix of incidence *X* for SNPs (TP), and *r* the vector of random residual effects assumed to be normally distributed 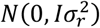.

We discarded markers with a minor allele frequency less than 5% (19,065 markers left), and tested different models typically used in genomic selection (Meuwissen et al. 2001; Habier et al. 2007; VanRaden 2008; Park and Casella 2008; Habier et al. 2011): Bayesian approach with a Gaussian prior for SNP effects, Bayesian approach with a scaled-t prior, Bayesian approach with a two component mixture prior with a point of mass at zero and a scaled-t slab, Bayesian approach with a two component mixture prior with a point of mass at zero and a Gaussian slab, and Bayesian approach with a Double-Exponential prior. These models are implemented in the R package `BGLR` v1.0.9 (Pérez and de los Campos 2014) available within `PopVar`, with the respective following notations: ‘BRR’ (Bayesian Ridge Regression), ‘BayesA’, ‘BayesB’, ‘BayesC’, and ‘BL’ (Bayesian Lasso).

We predicted the PM as the average parental GEBV computed as the matrix product between parental genotypes and estimated marker effects:

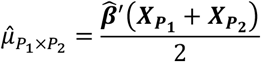

with 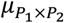 the PM for cross *P*_1_ × *P*_2_, 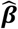 the vector of estimated marker effects, and 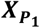 and 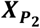 the vector of genotypes for parent *P*_1_ and parent *P*_2_ respectively.

For gametic variance, we considered the formula provided by Lehermeier et al. 2017:

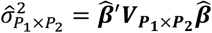

with 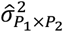 the gametic variance for cross *P*_1_ × *P*_2_, and 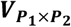 the genotypic variance-covariance matrix for biparental RIL progeny calculated as follows:

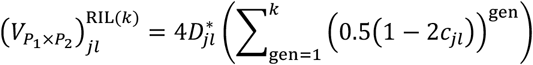

with 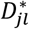 the Linkage Disequilibrium (LD) between alleles at *loci j* and *l* among both parental lines (either 0 if parents carry the same alleles at *loci j* and/or *l*, or 0.25 if alleles are different at both *locus* between parents and in coupling phase, that is one parent carries the two beneficial alleles while the other carries deleterious alleles, or - 0.25 if the alleles are in repulsion phase), *k* the number of generations, and *c*_*jl*_ the recombination rate between *loci j* and *l*. The recombination rates were computed from the West-European bread wheat population genetic map (Danguy des Déserts et al. 2021).

An alternative approach was used to compute the gametic variance with an in-house script. The model called Vg1 (standing for variance of gametes) is the RR-BLUP model (Meuwissen et al. 2001):

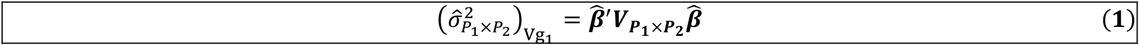

For the model called Vg2, we added a first term taking into account the error in marker effect estimation given by Lehermeier et al. 2017 (equation 10) in their algebraic version of the Posterior Mean Variance (PMV):

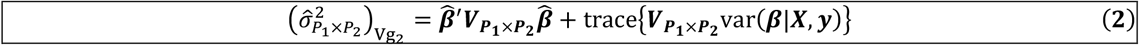

with 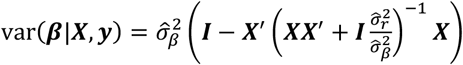, where 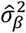 and 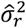 are the markers and residual estimated variances, and *X* is the vector of TP’s genotypes.

For the model called Vg3, we added a second term that aims to consider the fact that the uncertainty of the estimation of marker effects is modulated by the genomic constitution of each parent:

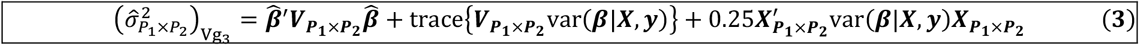

with 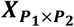 the vector of genotypes for the F1 of cross *P*_1_ × *P*_2_. Details are provided in **Supplementary Information S1**.

The UC was calculated as follows (Schnell and Utz 1975; Zhong and Jannink 2007):

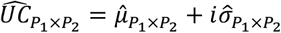

with *i* the selection intensity corresponding in our study to a 7% selection rate (*i* ∼ 1.91, computed as the inverse Mills ratio), and 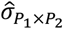 the estimated progeny SD for cross *P*_1_ × *P*_2_.

#### Experimental cross value components

We calculated the experimental cross value components for each cross from the distribution of the BLUE values: mean (PM), standard deviation (SD) and mean of the 7% best progeny (observed UC) (or the value of the best progeny when progeny size was inferior to 15).

#### Prediction ability

We compared PM, SD and UC genomic estimates with the corresponding experimental values using a weighted Pearson’s correlation (`weights` v1.0.4 R package (Pasek and Schwemmle 2021)) to adjust for the number of progenies per cross. These correlations were considered as prediction ability.

### Simulation study

In order to evaluate the different parameters impacting the prediction ability of the three cross value components (PM, SD and UC) and to have a reference for the values we can get in optimal/suboptimal conditions (perfect/unperfect prediction of marker effects, limited/infinite progeny size), we developed a simulation study that mimics our experimental design.

#### Simulation Design

The general scheme of the simulation study is provided in **Figure 1**. We based our simulations on real genotypes for the TP and the same parents’ genotypes for the crosses to be predicted. We also used in the formula the genetic map (Danguy des Déserts et al. 2021) estimated on real data for our predictions.

**Figure 1.**
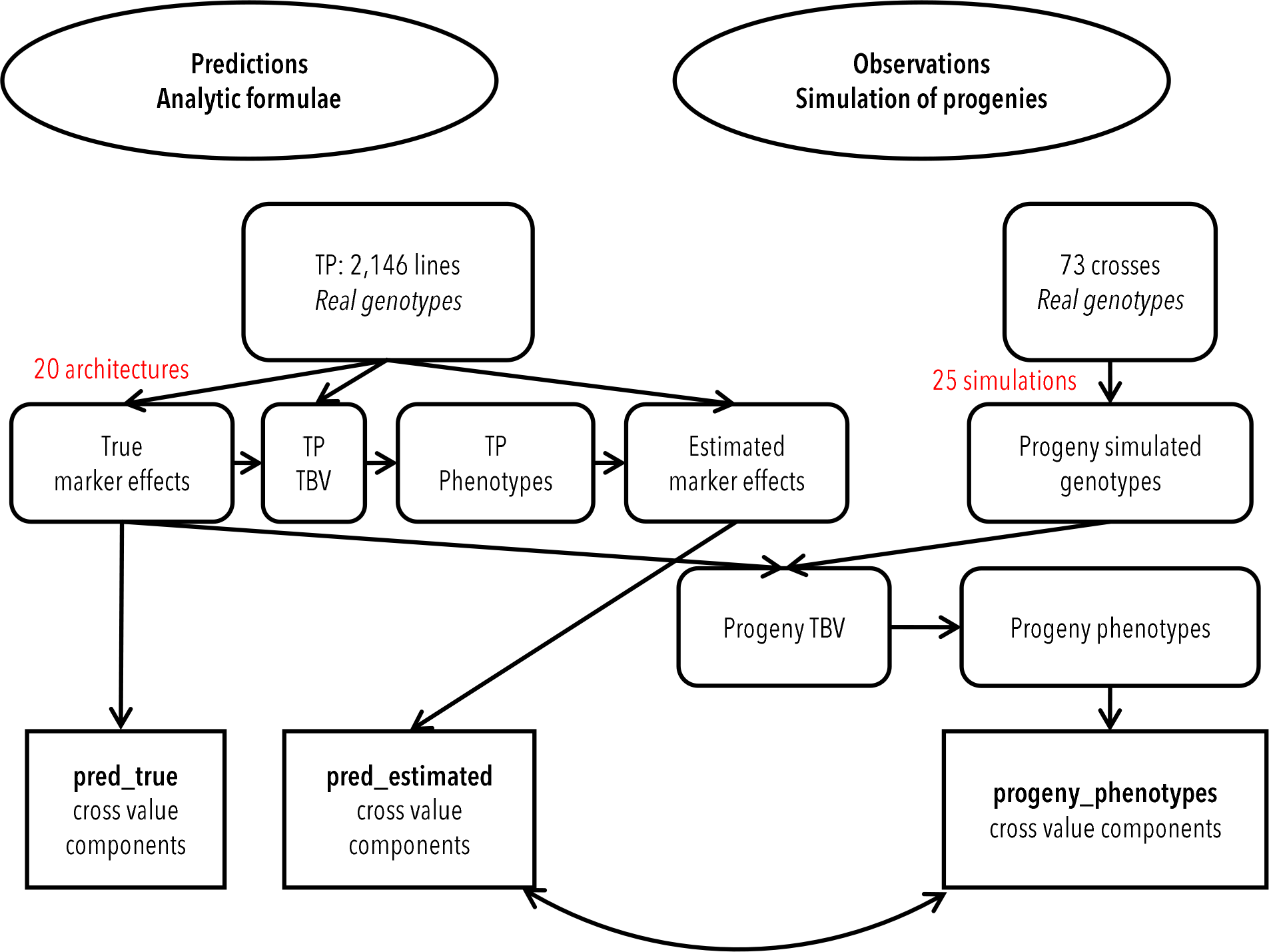
Simulation scheme. Abbreviations: TP: Training Population, TBV: True Breeding Value, pred_true: predicted true cross value components, pred_estimated: predicted estimated cross value components, progeny_phenotypes: observed (simulated phenotypes) cross value components.

### TP and predictions

On the one hand, the TP was composed of 2,591 genotyped varieties as described previously. We simulated Quantitative Trait *Loci* (QTL) by randomly selecting a specified number of markers (for instance 300) among the 21,340 markers (discarding the ones with a minor allele frequency less than 10%) and affect them a value drawn from the normal distribution *N*(0,1). These true marker effects were then adjusted to provide a variance of true breeding values (TBV) similar to the GEBV variance previously obtained in the experimental data for the yield trait. TBV were calculated as the cross product between true marker effects and genotypes. Next, we simulated phenotypes for the TP by adding a noise to their TBV. This noise was normally distributed with a variance corresponding to the residual variance from a specified heritability. For instance, if the TBV variance is 80 and we specify a heritability of 0.7, we would draw the noise for phenotypes computation from *N*(0,34) according to the following equation 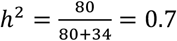

The 2,146 varieties that were phenotyped and genotyped (in the real TP) were used to estimate marker effects. True QTL markers were removed from this step. We tested different prediction models, as described previously, either using the different Bayesian models (`PopVar` R package), or using an in-house script implementing a RR-BLUP approach.

For each cross from our real experimental protocol, we computed the predictions for two scenarios: true and estimated PM, SD and UC (the three cross value components previously described) based on analytic formulae using true and estimated marker effects, respectively. The **pred_true** (**Figure 1**) scenario is the optimal situation where the sizes of the TP and of the progenies are infinite and where marker effects are perfectly estimated. In contrast, the **pred_estimated** (**Figure 1**) scenario corresponds to a situation where the size of the progenies is infinite but where marker effects are estimated using a prediction model in a fixed TP (similar scenario to our experimental protocol).

### Simulated progeny observations

We simulated progeny genotypes for the 73 crosses from our real experimental protocol (protein content phenotype for which we had the smallest number of crosses) using the `MoBPS` v1.6.64 R package (Pook et al. 2020). The numbers of progenies per cross were either the exact same values as in experimental data (**Supplementary Table S11**) or a fixed number for all crosses depending on the scenario (varying from 0 to 2,000, see next paragraph). We performed this step 25 times to account for the possibility that progeny genotypes might vary due to Mendelian gamete sampling (the results presented in this study are the average of these 25 simulations). Progeny TBV were calculated as the cross product between true simulated marker effects and the simulated progeny genotypes. Next, we simulated progeny phenotypes by adding a noise to their TBV. This noise was normally distributed with a variance corresponding to the residual variance observed in the TP (34 if we keep the example of the previous section with a heritability of 0.7).

Finally, we calculated the cross value components for each cross from the distribution of the progeny phenotypes (**progeny_phenotypes** in **Figure 1**): mean, standard deviation and mean of the 7% best progeny for the UC (or the value of the best progeny when progeny size was inferior to 15).

### Prediction ability

We compared the genomic predictions (true or estimated) with the simulated observations (true or phenotypes) using the weighted Pearson’s correlation to adjust for the number of progenies per cross. These correlations were considered as prediction ability.

#### Parameters influencing the prediction ability

For each scenario, 20 trait architectures were simulated. To compare scenarios, we considered for each cross value component the median values of correlations across the 20 runs.

We explored different parameters that could impact the prediction ability of the three cross value components: genetic architecture (number of QTL and heritability of the trait), number of progenies per cross, genomic prediction models, approach to estimate the gametic variance, and genetic constitution of the parents and their combination for crosses.

### Genetic architecture

We tested different number of QTL for the simulation of the genetic determinism of agronomic traits: 10-30-300-1,000. The scenarii with 10 or 30 QTL represent traits with major genes, whereas 300 and 1,000 QTL scenarii correspond to polygenic traits. Heritability ranged from 0.2 to 0.8 (with a 0.1 step). This heritability is the true heritability. To evaluate the quality of its estimation, we estimated in our simulations the heritability using a RR-BLUP approach implemented in the `rrBLUP` R package. As we observed that a true heritability of 0.7 corresponds to an estimated heritability of ∼0.5 in our simulations (**Supplementary Figure S1**), we considered 0.7 as a reference true heritability. It corresponds to a trait of 0.5 estimated heritability (yield trait for instance, see Results section).

### Number of progenies per cross

To evaluate if the observed variance in the field using a limited progeny size is reliable, we tested different numbers of progenies per cross, starting with the real numbers observed in the field (ranging from 5 to 122 progenies; mean = 47.6 progenies per cross, **Supplementary Table S11**) to a maximum of 2,000 progenies for all crosses (50, 100, 200, 1,000 and 2,000).

### Methods to estimate marker effects

To evaluate the importance of the quality of marker effects estimation in genomic predictions, we implemented a true and an estimated scenario. For the later, several genomic selection methods were compared: different Bayesian approaches and a RR-BLUP approach.

### Different approaches to estimate gametic variance

We implemented two algebraic variants of the PMV approach proposed by Lehermeier et al. 2017 with a RR-BLUP approach described earlier (‘Vg1’, ‘Vg2’, ‘Vg3’).

### Parents’ crosses

The experimental mating plan of the *PrediCropt* FSOV project was chosen by breeders to maximize genetic gain. To evaluate the bias possibly generated by expert choice (some parents were used in several crosses and some were genetically similar, as described in **Supplementary Table S11** and **Supplementary Figure S2**, respectively), we tested a different set of parents. This new set was chosen to be representative of the total genetic diversity present in the TP and the least possible genetically similar. We chose 146 different parents to generate 73 crosses. To do so, we first extracted the 10 Principal Components (PC analysis) from the variance-standardized Genomic Relationship Matrix (GRM) of the 2,591 genotyped varieties of the TP, using `PLINK` v2.00a2LM toolset. Next, we excluded varieties that did not have a phenotype for the yield trait in the real TP dataset (2,146). We computed euclidean distances on the 10 PC and performed a hierarchical cluster analysis (146 groups) with the Ward’s minimum variance method (Murtagh and Legendre 2014) (`dist` and `hclust` functions, respectively, from the `stats` v4.1.1 R package (R Core Team 2021)). In each cluster, we picked a line with the minimal value of the maximal relationship (GRM value) across all the remaining varieties, leading to the selection of 146 parents. Finally, we generated 73 crosses by randomly mating this new set of parents. This final procedure of random mating was replicated 10 times. We considered the median values of correlations across the 10 random matings to compare with the crosses from our experimental data. The 4 PC for the two sets of parents (“PrediCropt” crosses and this new set of “Unrelated” parents) are provided in **Supplementary Figure S2**.

## Results

### Quality of the Training Population

We used as TP the historical French registration data (GEVES) and the INRAE-AO breeding program, both Southern and Northern France networks. We checked the repeatability of observations in the TP by computing Pearson’s correlations between years for each trait and each dataset (**Supplementary Tables S2** to **S9**). The average correlations per dataset were relatively high and ranged from 0.59 (INRAE-AO, South) to 0.81 (GEVES, South), with a mean of 0.69 all dataset combined for yield, from 0.65 (INRAE-AO, South) to 0.85 (GEVES, South), with a mean of 0.78 for grain protein content, from 0.81 (INRAE-AO, South) to 0.92 (GEVES, North), with a mean of 0.87 for plant height, and from 0.86 (INRAE-AO, South) to 0.96 (GEVES, North), with a mean of 0.91 for heading date.

The variance components, phenotypic repeatabilities and genomic prediction accuracies of the TP are provided in **Table 1**. The repeatabilities at the plot level were relatively low for yield (0.32), moderate for grain protein content (0.52) and high for plant height and heading date (0.73 and 0.85, respectively). Repeatabilities at the design level were very high for all traits (ranging from 0.91 to 0.99). All these values reinforce the good correlations obtained above and suggest a high-quality TP dataset. The genomic heritabilities were moderate for yield (0.53), grain protein content (0.51) and plant height (0.56), and high for heading date (0.73). Finally, the cross-validation accuracy was quite similar for all traits, with moderate to high values ranging from 0.52 (plant height) to 0.67 (heading date).

**Table 1.** Variance components, repeatabilities and cross-validation accuracy in the TP. 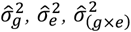 and 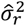 are the line, the environment, the interaction between line and environment, and the residual estimated variances using a mixed model with trials as random effect. 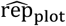 and 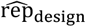 are repeatabilities at the plot and design levels, computed from these estimated variances. 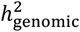 was estimated using RR-BLUP model. The cross-validation accuracy was estimated using 60% of the data as the training population, and 40% as the validation population.

| Trait | $\hat{\sigma}_g^2$ | $\hat{\sigma}_e^2$ | $\hat{\sigma}_{(g \times e)}^2$ | $\hat{\sigma}_r^2$ | $\widehat{\text{rep}}_{\text{plot}}$ | $\widehat{\text{rep}}_{\text{design}}$ | $\hat{h}_{\text{genomic}}^2$ | Cross-validation accuracy |
| --- | --- | --- | --- | --- | --- | --- | --- | --- |
| Yield | 18.9 | 237.1 | 24.8 | 15.4 | 0.32 | 0.91 | 0.53 | 0.66 ($\pm 0.018$) |
| Grain protein content | 0.3 | 1.0 | 0.2 | 0.1 | 0.52 | 0.93 | 0.51 | 0.60 ($\pm 0.018$) |
| Plant height | 32.6 | 56.5 | 6.5 | 5.6 | 0.73 | 0.97 | 0.56 | 0.52 ($\pm 0.021$) |
| Heading date | 12.6 | 65.5 | 1.9 | 0.5 | 0.85 | 0.99 | 0.73 | 0.67 ($\pm 0.016$) |

### Simulation study

#### Genetic architecture

In our simulation study, we explored different scenarii for the genetic architecture in order to see its impact on the prediction ability of the three cross value components (PM, SD and UC).

The prediction ability of the three cross value components decreased when QTL number increased (**Figure 2**). PM and UC were moderately affected: the prediction abilities decreased from 0.91 to 0.84 and from 0.82 to 0.77 between 10 and 1,000 QTL (heritability = 0.6), for PM and UC respectively. SD was severely affected with prediction abilities dropping from 0.69 to 0.18 between 10 and 1,000 QTL for the same scenario.

**Figure 2.**
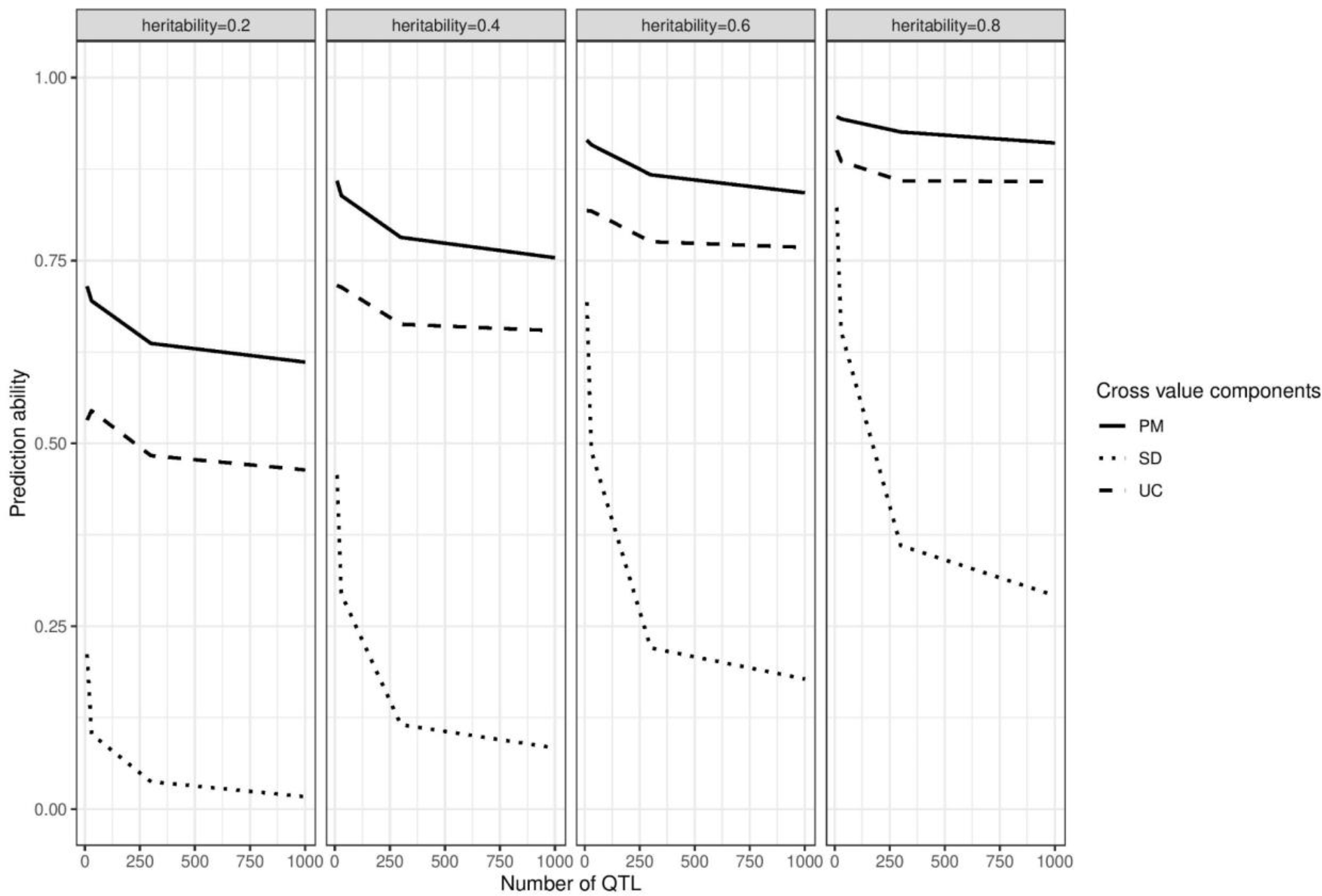
Impact of genetic architecture on the prediction ability of the three cross value components, based on simulations. The number of progenies per cross was the same as in experimental data and the marker effects were estimated with ‘BayesA’ method. Abbreviations: PM: Parental Mean, SD: Standard Deviation, UC: Usefulness Criterion.

The prediction ability of the three cross value components increased when heritability increased (**Figure 2**). The impact was quite similar for the three components: the prediction abilities increased from 0.64 to 0.93, from 0.48 to 0.86, and from 0.04 to 0.36 between heritability 0.2 and heritability 0.8 (number of QTL = 300), for PM, UC and SD respectively.

#### Progeny size

To evaluate if the observed variance in the field using a limited progeny size (< 100) is reliable, we tested different numbers of progenies per cross (**Figure 3**).

**Figure 3.**
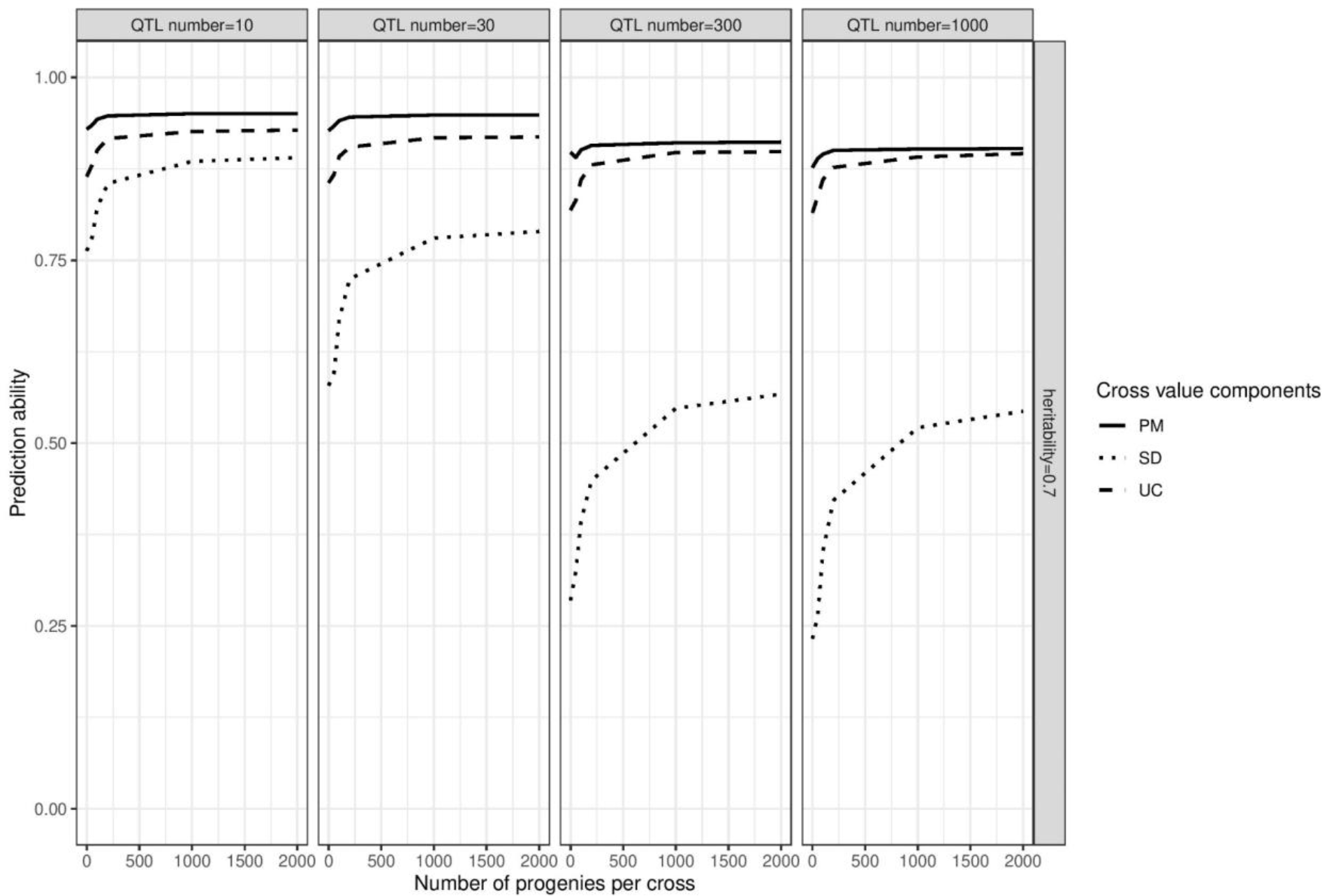
Impact of the number of progenies per cross on the prediction ability of the three cross value components, based on simulations. Marker effects were estimated with ‘BayesA’ method. Abbreviations: PM: Parental Mean, SD: Standard Deviation, UC: Usefulness Criterion.

The prediction ability of the three cross value components increased when the number of progenies per cross increased (**Figure 3**). PM and UC were moderately affected: the prediction abilities increased from 0.90 to 0.91 and from 0.82 to 0.90 between experimental number of progenies per cross and 2,000 progenies per cross (heritability = 0.7 and QTL number = 300), for PM and UC respectively. SD was severely affected with prediction abilities increasing from 0.29 to 0.57 between experimental number of progenies per cross and 2,000 progenies per cross for the same scenario. We observe a first bow for 250 progenies and a second for 1000, close to a plateau.

The relative increase of prediction ability for SD was remarkably stronger in scenarii with high number of QTL (**Figure 3** and **Supplementary Figure S3**) although there is little improvement between the 300 and 1,000 QTL scenarii.

#### Knowledge of QTL

Genomic predictions are based on the hypothesis that we are able to estimate QTL effects using a representative TP. We used as a reference a scenario (**pred_true** in **Figure 1**) which is the optimal situation with infinite TP size and where marker effects are perfectly estimated (we use true simulated marker effects to estimate PM, SD and UC). In contrast, the **pred_estimated** (**Figure 1**) scenario corresponds to a situation where marker effects are estimated using a prediction model in a limited TP. These two scenarii are provided in **Figure 4 A**.

**Figure 4.**
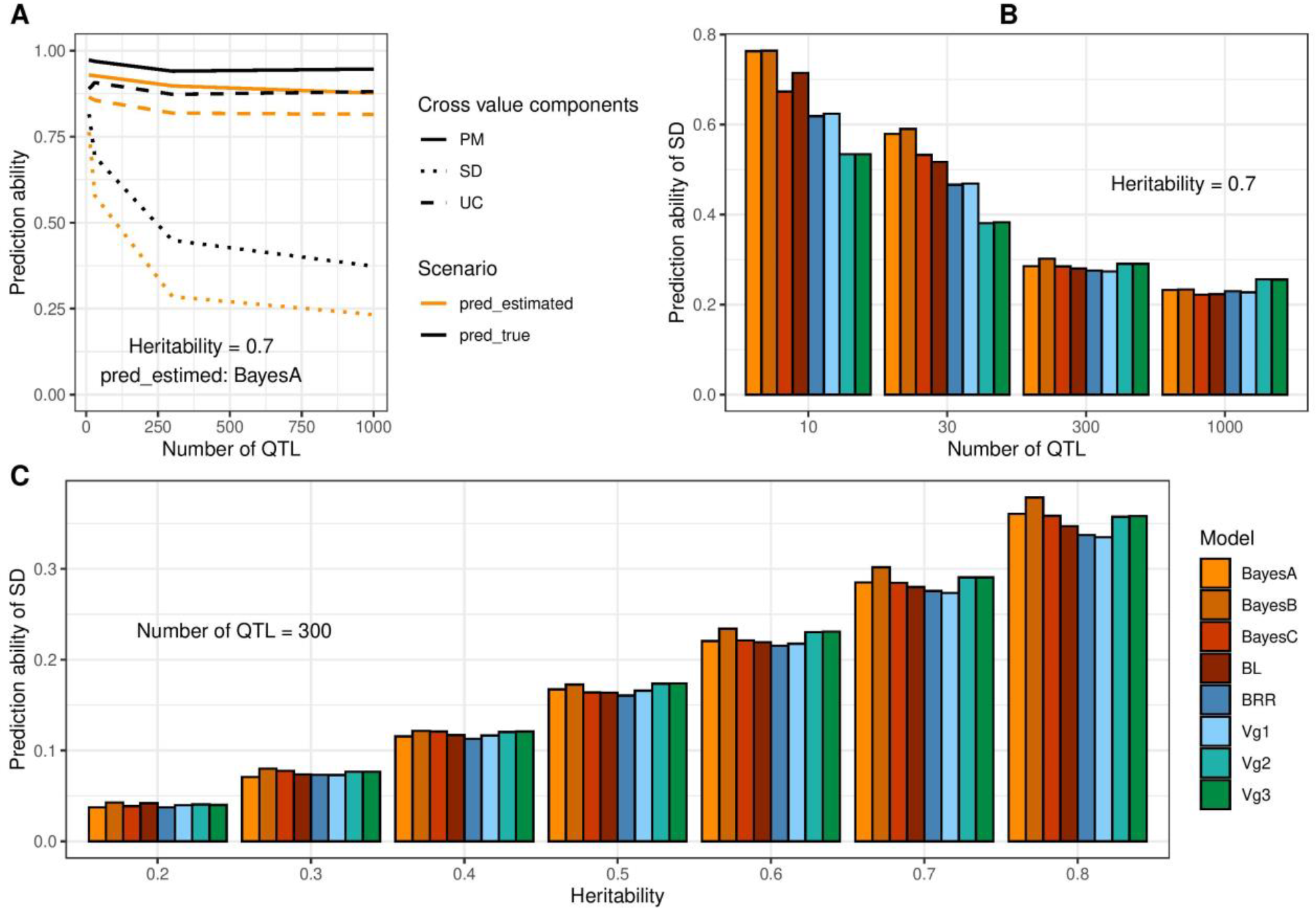
Impact of the knowledge of QTL on the prediction ability of the three cross value components (A) and of the genomic selection method on the prediction ability of SD (B and C), based on simulations. The number of progenies per cross was the same as in experimental data. In A, the heritability was 0.7 and marker effects were estimated with ‘BayesA’ method. The true cross value components (pred_true) were calculated with true marker effects whereas the estimated cross value components (pred_estimated) were calculated with estimated marker effects. In B, the heritability was 0.7. In C, the number of QTL was 300. Abbreviations: PM: Parental Mean, SD: Standard Deviation, UC: Usefulness Criterion.

In the TRUE scenario (**pred_true**), the prediction abilities were very high for the three cross value components with a low QTL number (10): 0.97, 0.89 and 0.81 for PM, UC and SD, respectively (**Figure 4 A**) when the number of progenies was the same as in the experimental data (0 in Figures), and 0.99, 0.95 and 0.95 when the number of progenies was 2,000 (**Supplementary Figure S4**). We observed a strong decrease of prediction ability for SD when the number of QTL increased (0.37 for QTL number = 1,000 *versus* 0.81 for QTL number = 10, **Figure 4 A**) when progeny was inferior to 100 (experimental design), and much smaller when the number of progenies per cross was 2,000 (0.86 for QTL number = 1,000 *versus* 0.95 for QTL number = 10, **Supplementary Figure S4**). All these results show that if we perfectly know the QTL effects, we are able to very well estimate the three cross value components.

As we observed an important gap of prediction abilities between the **pred_true** and the **pred_estimated** scenarii for SD (**Figure 4 A** and **Supplementary Figure S4 A**), we tested different prediction models with different hypotheses on trait architecture. We also tested if taking into account explicitly in the analytic formulae of SD the error of estimation of marker effects had a significant impact (see Materials and methods). In total, we compared 6 prediction models: BayesA, BayesB, BayesC, BL, BRR, and RR-BLUP (Vg1), and 3 SD estimators: Vg1 (RR-BLUP), Vg2 (Vg1 + 1 term), and Vg3 (Vg2 + 1 term) (**Figure 4 B & C**).

Looking at small progeny size (equal to the experimental design) scenarii, with low number of QTL (QTL number = 10 and heritability = 0.7; **Figure 4 B**), Bayesian models performed better than the RR-BLUP approach, and more particularly the Bayes A and B models (prediction abilities of 0.76 for both Bayes A and B models *versus* 0.62 for Vg1). As seen previously, little difference was observed between 300 QTL and 1,000 QTL scenarii. With 300 QTL, the differences of prediction abilities between models were small, ranging from 0.27 (Vg1) to 0.30 (BayesB). Taking into account the error in marker effect estimates slightly increased the prediction abilities in scenarii with high number of QTL (0.29 for Vg2 and Vg3 with 300 QTL). The ranking of prediction models for 300 QTL scenarii was the same for increasing heritabilities (**Figure 4 C**).

Looking at large progeny size scenarii (2,000 progenies per cross), the differences of prediction abilities for SD between models were slightly increased (0.91 for Bayes B *versus* 0.73 for Vg1 in QTL number = 10 and heritability = 0.7 scenario; **Supplementary Figure S4 B**). Vg3 increased the prediction abilities, especially in low heritability scenarii (0.25 for Vg1 *versus* 0.33 for Vg3 in QTL number = 300 and heritability = 0.2 scenario; **Supplementary Figure S4 C**).

All these results show that the quality of marker effect estimates has a strong impact on the prediction ability of SD. For traits with small number of QTL, Bayesian models (Bayes A or B) should be preferred. For traits with high number of QTL, when heritability is high RR-BLUP is fine, when heritability is low Vg3 should be used.

#### Crosses selected for predictions

To evaluate the bias possibly generated by the choice of crosses in our experimental design, we compared the results obtained with the experimental crosses called “PrediCropt” with a set of crosses called “Unrelated”. We selected the “Unrelated” parents to be representative of the total genetic diversity present in the TP and the crosses by mating parents with low genetic similarity (see Materials and methods). The 4 PC for the two sets of parents are provided in **Supplementary Figure S2** and show that the “Unrelated” parents are more scattered throughout the TP than “PrediCropt” parents (for instance, no “PrediCropt” parent in the bottom left side in graphic **A** - PC 1 *versus* PC 2 -).

Ten random associations of the “Unrelated” set of parents gave the prediction abilities’ results provided in **Figure 5** for scenario with 300 QTL and heritability of 0.7. Little differences were observed between the two sets of parents. Indeed, the prediction abilities were 0.90 (“PrediCropt”) *versus* 0.88 (“Unrelated”) for PM, 0.82 for both sets for UC, and 0.29 (“PrediCropt”) *versus* 0.24 (“Unrelated”) for SD. According to these results, we considered as negligible the bias generated by the choice of our experimental crosses.

**Figure 5.**
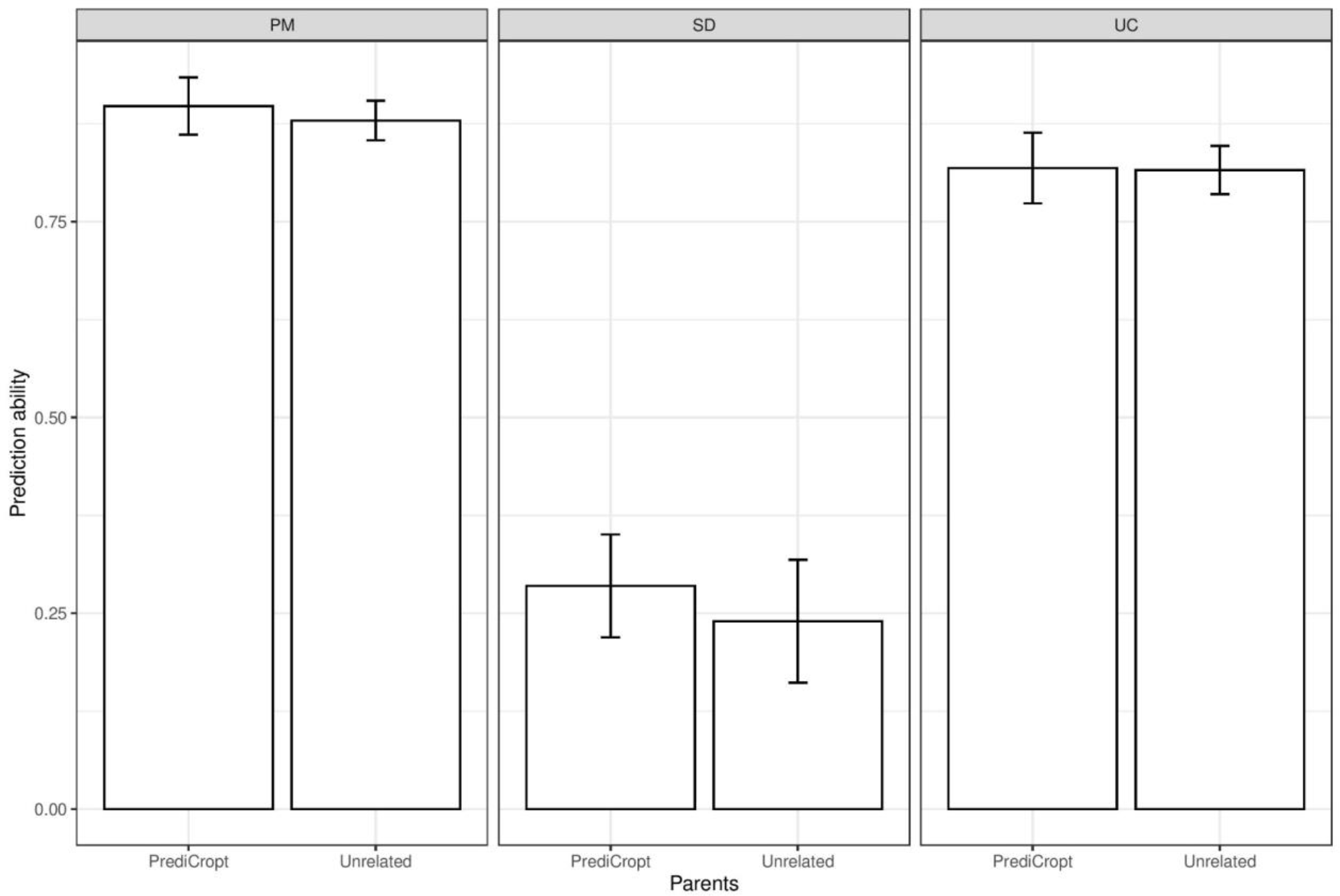
Impact of the genetic constitution of the crosses’ parents on the prediction ability of the three cross value components, based on simulations. “PrediCropt” parents are the same crosses as in experimental data whereas “Unrelated” parents are 10 random associations of a set of unrelated parents selected as described in the Materials and methods section ((median values given). The number of progenies per cross was the same as in experimental data. The heritability was 0.7 and the number of QTL, 300. Marker effects were estimated with ‘BayesA’ method. Abbreviations: PM: Parental Mean, SD: Standard Deviation, UC: Usefulness Criterion.

### Experimental data application

In our experimental data, we phenotyped 4 traits of the progenies of the 101 or 73 PrediCropt crosses depending on the trait (**Supplementary Table S11**). In order to check the reliability and consistency of phenotypes between sites and years, we computed Pearson’s correlations (**Supplementary Tables S12 and S13**). On average across all sites and years, correlations were 0.48, 0.43, 0.79 and 0.88 for yield, grain protein content, plant height, and heading date, which were quite good values for such an experimental design.

We compared experimental values to the genomic predictions computed using the TP described earlier. The prediction abilities of the three cross value components in experimental data are provided in **Table 2**. PM was correctly predicted for the 4 phenotypes (median values across all prediction models: 0.38, 0.63, 0.51 and 0.91, for yield, grain protein content and heading date, respectively). SD was correctly predicted for plant height and heading date (median values: 0.59 and 0.38, respectively), and badly predicted for yield and grain protein content (median values: 0.01 and 0.13, respectively). Finally, UC was correctly predicted for all 4 traits (median values: 0.45, 0.52, 0.54 and 0.74 for yield, grain protein content, plant height and heading date, respectively).

**Table 2.** Prediction ability (PA) of the three cross value components in experimental data. Prediction ability was computed as weighted Pearson’s correlation between genomic predictions and phenotype observations. Prediction models are the following: BayesA (Bayesian approach with a scaled-t prior for SNP effects), BayesB (Bayesian approach with a two component mixture prior with a point of mass at zero and a scaled-t slab), BayesC (Bayesian approach with a two component mixture prior with a point of mass at zero and a Gaussian slab), BL - Bayesian Lasso - (Bayesian approach with a Double-Exponential prior), BRR - Bayesian Ridge Regression - (Bayesian approach with a Gaussian prior), and Vg - Variance of gametes - (RR-BLUP approach with a Gaussian distribution for SNP effects): 1 using the same formula for calculating gametic variance as in the previous models, 2 and 3 are two algebraic variants of the PMV approach proposed by Lehermeier et al. 2017 (see Materials and methods). Abbreviations: PA: Prediction Ability, CI: Confidence Interval, PM: parental mean, SD: standard deviation, UC: usefulness criterion.

| Trait | Model | PM |  | SD |  | UC |  |
| --- | --- | --- | --- | --- | --- | --- | --- |
|  |  | PA | CI | PA | CI | PA | CI |
| Yield | BayesA | 0.37 | 0.18, 0.55 | 0.02 | -0.18, 0.22 | 0.45 | 0.28, 0.63 |
|  | BayesB | 0.38 | 0.19, 0.56 | 0.05 | -0.15, 0.24 | 0.47 | 0.29, 0.64 |
|  | BayesC | 0.38 | 0.19, 0.56 | 0.02 | -0.18, 0.21 | 0.46 | 0.28, 0.63 |
|  | BL | 0.38 | 0.20, 0.56 | 0.01 | -0.19, 0.21 | 0.45 | 0.28, 0.63 |
|  | BRR | 0.38 | 0.20, 0.56 | 0.00 | -0.20, 0.20 | 0.45 | 0.28, 0.63 |
|  | Vg1 | 0.38 | 0.20, 0.56 | -0.01 | -0.20, 0.19 | 0.46 | 0.28, 0.63 |
|  | Vg2 |  |  | -0.17 | -0.36, 0.02 | 0.45 | 0.27, 0.62 |
|  | Vg3 |  |  | -0.17 | -0.36, 0.02 | 0.45 | 0.27, 0.62 |
| Grain protein content | BayesA | 0.63 | 0.44, 0.81 | 0.17 | -0.05, 0.40 | 0.52 | 0.32, 0.72 |
|  | BayesB | 0.62 | 0.44, 0.80 | 0.26 | 0.03, 0.48 | 0.52 | 0.33, 0.72 |
|  | BayesC | 0.63 | 0.45, 0.81 | 0.07 | -0.16, 0.30 | 0.51 | 0.31, 0.71 |
|  | BL | 0.63 | 0.45, 0.81 | 0.10 | -0.14, 0.33 | 0.51 | 0.31, 0.71 |
|  | BRR | 0.63 | 0.45, 0.81 | 0.08 | -0.15, 0.32 | 0.51 | 0.31, 0.71 |
|  | Vg1 | 0.63 | 0.45, 0.81 | 0.05 | -0.19, 0.28 | 0.52 | 0.32, 0.72 |
|  | Vg2 |  |  | 0.16 | -0.07, 0.39 | 0.53 | 0.33, 0.72 |
|  | Vg3 |  |  | 0.16 | -0.07, 0.39 | 0.53 | 0.33, 0.72 |
| Plant height | BayesA | 0.51 | 0.34, 0.68 | 0.64 | 0.48, 0.79 | 0.66 | 0.52, 0.81 |
|  | BayesB | 0.51 | 0.34, 0.68 | 0.63 | 0.48, 0.79 | 0.66 | 0.51, 0.81 |
|  | BayesC | 0.51 | 0.34, 0.68 | 0.64 | 0.49, 0.79 | 0.65 | 0.50, 0.80 |
|  | BL | 0.51 | 0.34, 0.68 | 0.64 | 0.49, 0.79 | 0.60 | 0.44, 0.75 |
|  | BRR | 0.50 | 0.32, 0.67 | 0.53 | 0.36, 0.70 | 0.47 | 0.30, 0.64 |
|  | Vg1 | 0.50 | 0.33, 0.67 | 0.54 | 0.37, 0.70 | 0.48 | 0.31, 0.65 |
|  | Vg2 |  |  | 0.41 | 0.23, 0.59 | 0.45 | 0.27, 0.62 |
|  | Vg3 |  |  | 0.41 | 0.23, 0.59 | 0.45 | 0.27, 0.62 |
| Heading date | BayesA | 0.92 | 0.85, 1.00 | 0.46 | 0.28, 0.63 | 0.70 | 0.56, 0.84 |
|  | BayesB | 0.93 | 0.85, 1.00 | 0.46 | 0.29, 0.64 | 0.71 | 0.57, 0.85 |
|  | BayesC | 0.82 | 0.71, 0.93 | 0.49 | 0.32, 0.66 | 0.64 | 0.49, 0.79 |
|  | BL | 0.92 | 0.84, 1.00 | 0.41 | 0.23, 0.59 | 0.71 | 0.58, 0.85 |
|  | BRR | 0.90 | 0.82, 0.99 | 0.33 | 0.14, 0.52 | 0.77 | 0.64, 0.90 |
|  | Vg1 | 0.90 | 0.82, 0.99 | 0.34 | 0.16, 0.53 | 0.77 | 0.64, 0.90 |
|  | Vg2 |  |  | 0.33 | 0.14, 0.51 | 0.78 | 0.65, 0.90 |
|  | Vg3 |  |  | 0.33 | 0.14, 0.51 | 0.78 | 0.65, 0.90 |

As expected according to the results of our simulation study, little differences were observed between prediction models (**Table 2**). One notable result was for plant height: SD was remarkably better predicted with non-Gaussian Bayesian approaches than with Gaussian approaches (prediction ability of 0.63-0.64 with non-Gaussian Bayesian approaches *versus* 0.53-0.54 with Gaussian approaches), suggesting that this trait is determined by few QTL according to the results of our simulation study.

## Discussion

The present study aimed at investigating the genomic prediction ability of the cross value called UC and its two components, the mean (PM) and the variance (SD) of the progeny. We used simulations to test the impact of the error of marker effect estimation in predictions: we compared two scenarii, one using true marker effects and one using estimated marker effects. We also tested the impact of trait heritability (varying from 0.2 to 0.8), trait architecture (number of QTL varying between 10 and 1,000) and progeny size (varying from the number of progenies per cross observed in the field to 2,000). Those figures served as a reference grid for the comparison of predictions with experimental data composed of 73 crosses with 60 progenies on average. Our results shed light on the factors influencing cross value prediction ability and provide insights into the prerequisites for using such methods in precision breeding programs.

### A high-quality TP

In this study, we took advantage of a historical TP of more than 20 years of data to predict three cross value components (PM, SD and UC) of 73 or 101 crosses depending on the trait using genomic prediction models. We compared these predictions to experimental observations of progenies. All the results concerning the TP (correlations between years, repeatabilities at plot and design levels) suggest a high-quality dataset. The prediction accuracies (cross-validation) ranging from 0.52 (plant height) to 0.67 (heading date) were higher than previously obtained with a smaller TP (G. Charmet, personal communication). Moreover, based on preliminary studies (data not shown) inspired from an iterative approach that removes the environments (year x location) one by one and that computes prediction accuracy at each iteration (Heslot et al. 2013), we kept all the environments of the TP because we did not detect any environment deteriorating the quality of the prediction. We also checked (data not shown) the prediction accuracy within each sub-dataset of the TP (GEVES, INRAE-AO, North, South) and the prediction abilities of the 73 crosses obtained for the three cross-value components using these sub-datasets as TP. We observed better prediction abilities using all datasets combined. All these preliminary results led us to retain the entire dataset as the TP in this study.

Due to the size and complexity of the TP design, for computational reasons (the need of pre-corrected phenotypes in the R package, and time to run), and as done in previous studies (Tiede et al. 2015; Danguy des Déserts et al. 2023), we separated the steps of correction for the fixed effects such as spatial coordinates and environmental effects (location and year) from the estimation of marker effects. It would be interesting to evaluate the impact of this pre-correction on the estimation of marker effects step.

### Parameters impacting prediction abilities based on simulations

To study the conditions necessary to obtain good prediction abilities in our experimental design, we simulated progenies for our crosses in such a way as to mimic this design, fixing parameters such as heritability or number of QTL to generate different trait architectures based on the real TP genotypes.

The first comment on this simulation study concerns the difference observed between true heritability and estimated heritability. We observed that we lost between 0.1 and 0.2 between true and estimated heritabilities, whatever the number of QTL simulated. This missing heritability, already reported before, could be explained by a lack of genetic information, such as a linkage disequilibrium not perfectly known in our genotype set, potentially due to the absence of rare variants or structural variants (Manolio et al. 2009), epistasis and GxE. According to this observation, we focused our results on scenarii where heritability = 0.7, which correspond to an estimated heritability of around 0.5 to match with the estimations for the traits in our experimental design.

The first parameter having a very strong impact on the prediction abilities was the heritability: the prediction ability of the three cross value components increased when heritability increased. The impact of QTL number was weaker except for SD prediction: the prediction ability decreased when QTL number increased. These two results were in concordance with the results obtained in previous studies (Wimmer et al. 2013; Tiede et al. 2015; Yao et al. 2018). The first conclusion about this simulation study is that trait architecture is the most important factor impacting prediction abilities and more especially for the prediction of SD.

A second parameter strongly impacting the quality of prediction was the progeny size. The prediction ability of the three cross value components increased when the number of progenies per cross increased, the SD being strongly affected. To our knowledge, no study has yet investigated this parameter. Our results suggest that more than 1,000 progenies per cross should be sufficient to maximize SD prediction ability as little differences were observed between the 1,000 and 2,000 progenies per cross scenarii.

We investigated another parameter: the quality of estimation of marker effect. We observed that if we perfectly knew marker effects (TRUE scenario), we were able to almost perfectly estimate the three cross value components (0.99, 0.95 and 0.95 for PM, UC and SD respectively in scenario with heritability = 0.7, QTL number = 10, and 2,000 progenies per cross). The formulae for predictions are thus correct and it seems that we lose information at the step of estimating marker effects. This could be due to two factors: a TP not informative enough compared to our validation population, or/and a prediction model not efficient. We could not explore the first factor because no other TP was available for this study. However, we investigated the second parameter by testing different models of prediction. We observed small differences between prediction models, and mostly in the progeny size scenarii. This result was in concordance with several previous studies (Yao et al. 2018; Neyhart et al. 2019; Santos et al. 2019). Although RR-BLUP is known to have a similar or slightly better performance for the GEBV prediction than differential shrinkage models for scenarii with a large number of QTL (Daetwyler et al. 2010), we were expecting better accuracies of the estimated marker effects using differential shrinkage models for scenarios with a low number of QTL (Meuwissen et al. 2001; Shepherd et al. 2010; Legarra et al. 2011). We indeed observed this result: with low number of QTL (10, 30), Bayesian models performed slightly better than the RR-BLUP approach, and more particularly the Bayes A and B models.

In this study, we computed SD using the algebraic formula proposed by Lehermeier et al. 2017 for gametic variance. As the step of estimating marker effects was reducing prediction abilities, we implemented two variants for the computation of SD to account for the error in marker effect estimation. The first variant (Vg2) was the algebraic version of the Posterior Mean Variance (PMV) (Lehermeier et al. 2017). The second variant (Vg3) aimed at considering the fact that the uncertainty of the estimation of marker effects is modulated by the genomic constitution of each parent. These two variants were computed only with marker effects estimated by a RR-BLUP approach in our study. They increased SD prediction abilities with high number of QTL and to a greater extent with low heritability. This result is in concordance with the 300 QTL simulation scenario described in Lehermeier et al. 2017.

This simulation study considered a fixed set of scenarios, and further investigations into a broader range of genetic architectures and trait complexities would be beneficial. Notably, this study could be improved by modeling traits with an interaction between genotype and environment parameter, which could be a closer scenario to reality.

### Validation in experimental data

To our knowledge, this study is the first study that evaluates the prediction ability of cross value components based on real progenies’ phenotypes observed in experimental data. The prediction abilities of PM and UC were good for the 4 traits. The prediction abilities of SD varied depending on the trait and the choice of prediction models. For instance, we were not able to significantly predict the SD for yield. Yield is a very complex trait probably influenced by environmental effects, as seen from the results of the variance components estimation, which could make the variability of the phenotype difficult to predict. For the 3 other traits, non-Gaussian Bayesian approaches outperformed Gaussian approaches for SD prediction, indicating that considering more flexible distributions for marker effects can improve SD estimation accuracy. This observation is particularly relevant for plant height, where SD prediction exhibited significant improvements with non-Gaussian models. This suggests that the underlying genetic architecture of plant height might be determined by a few major genes (Ellis et al. 2002), which are better captured by non-Gaussian approaches.

While this experimental data application demonstrated promising results, more extensive validation across diverse datasets and crops is warranted to validate the generalizability of our findings. In particular, it would be interesting to generate more progenies per cross since the simulation results showed that with more progenies per cross, prediction abilities were much better. It also would be interesting to validate the interest of criteria such as UC in more diverse material, in a pre-breeding program for instance. Finally, as the negative genetic correlation between yield and grain protein content is a highly interesting issue for the wheat industry (Thorwarth et al. 2018), one could also study the prediction ability of this correlation using the predictive analytical formulae that have been recently developed (Neyhart et al. 2019).

### Recommendations for breeding schemes

Our results contribute to the growing body of knowledge on genomic selection and provide practical guidance for precision breeding programs. The study highlights the importance of accurate marker effect estimates, gathering of large and diverse TP datasets, and the selection of appropriate prediction models for improving cross value components prediction abilities. The findings can aid breeders in prioritizing resources and designing more efficient breeding strategies. For instance, according to our results, it would be promising to generate lots of progenies for the most promising crosses according to UC, in order to maximize the chance to have the outstanding expected line according to predictions. Differential progeny sizes according to UC thresholds should be tuned by each breeder according to their genetic material and budget.

## Acknowledgements

The authors thank the *Fonds de Soutien à l’Obtention Végétale* (FSOV) for financing the FSOV 2020 I - PrediCropt project.

The authors acknowledge the partners of the project (Agri-Obtentions and Florimond-Desprez veuve & fils) and the INRAE personal responsible for experimental evaluation (Laurent Falchetto, Sandrine Berges -INRAE UE PHACC-, Clement Debiton -INRAE UMR GDEC, CRB group-, Kevin Bargoin -INRAE UMR GDEC, DIGEN group-, Patrice Walczak -INRAE UE Ferlus-, Paul Bataillon -INRAE UE Auzeville-, and Emmanuel Heumez - INRAE UE GCIE-). They also thank Marie-Hélène Bernicot and Solène Barrais for providing the GEVES training population dataset (corresponding to the experimental evaluation data for French national registration).

## Statements and Declarations

## Funding

This work was supported by the *Fonds de Soutien à l’Obtention Végétale* (FSOV 2020 I - PrediCropt project). Genotyping was supported by the Breedwheat grant (ANR-10-BTBR-0003) and INRAE IVD program.

## Competing Interests

The authors have no relevant financial or non-financial interests to disclose.

## Author Contributions

COE collected and formatted the data, performed the analyses, interpreted the results, and wrote the manuscript. EH, LD, EGD and FXO provided crosses’ progenies for the FSOV *PrediCropt* project. SB conceived and coordinated the FSOV project. JME and SB supervised COE, interpreted the results and helped writing the manuscript. All authors reviewed and approved the final version of the manuscript.

## Data availability

The data that support the findings of this study are openly available in repository FSOV PrediCropt at https://doi.org/10.57745/F6230V, reference number F6230V.

## Ethics approval

This is an observational study. No ethical approval is required.

## Notes

### Competing Interest Statement

The authors have declared no competing interest.

