## Supplementary for "Validation of cross progeny variance genomic prediction using simulations and experimental data in winter elite bread wheat"

**Supplementary Information S1. Algebric Posterior Mean Variance (PMV) approach for gametic variance computation.**

*Estimation of marker effects:*

We consider the following SNP-BLUP model to estimate marker effects in a Training Population (TP):

$$\boldsymbol{y} = \boldsymbol{\mu} + \boldsymbol{X\beta} + \boldsymbol{r}$$

with $\boldsymbol{y}$ the vector of phenotypes of the TP corrected for any fixed effects, $\boldsymbol{\mu}$ the grand mean, $\boldsymbol{\beta}$ a vector of random SNP effects assumed to be normally distributed $N\left(0, \boldsymbol{I}\sigma_\beta^2\right)$, with its matrix of incidence $\boldsymbol{X}$ for SNPs (TP), and $\boldsymbol{r}$ the vector of random residual effects assumed to be normally distributed $N(0, I\sigma_r^2)$.

We can develop the estimates of marker effects as:

$$\widehat{\boldsymbol{\beta}} = \mathrm{E}[\boldsymbol{\beta}|\text{data}] = \mathrm{E}[\boldsymbol{\beta}|\boldsymbol{X},\boldsymbol{y}] = \mathrm{E}[\boldsymbol{\beta}] + \mathrm{cov}(\boldsymbol{\beta},\boldsymbol{y})\mathrm{var}(\boldsymbol{y})^{-1}(\boldsymbol{y} - \widehat{\boldsymbol{\mu}})$$

with $\mathrm{cov}(\boldsymbol{\beta},\boldsymbol{y}) = \mathrm{cov}(\boldsymbol{\beta},\boldsymbol{X\beta}) = \hat{\sigma}_\beta^2\boldsymbol{X}$ and $\mathrm{var}(\boldsymbol{y}) = \boldsymbol{X}\hat{\sigma}_\beta^2\boldsymbol{X}' + \hat{\sigma}_r^2$, that leads to:

$$\widehat{\boldsymbol{\beta}} = \boldsymbol{X}'(\boldsymbol{XX}' + \boldsymbol{I}\theta)^{-1}(\boldsymbol{y} - \widehat{\boldsymbol{\mu}}) \qquad (1)$$

with $\theta = \frac{\hat{\sigma}_r^2}{\hat{\sigma}_\beta^2}$.

We can then deduce:

$$\mathrm{var}(\boldsymbol{\beta}|\text{data}) = \mathrm{var}(\boldsymbol{\beta}) - \mathrm{cov}(\boldsymbol{\beta},\boldsymbol{y})\mathrm{var}(\boldsymbol{y})^{-1}\mathrm{cov}(\boldsymbol{y},\boldsymbol{\beta}) = \sigma_\beta^2\boldsymbol{I} - \sigma_\beta^2\sigma_\beta^2\boldsymbol{X}'\left(\boldsymbol{XX}'\hat{\sigma}_\beta^2 + \boldsymbol{I}\hat{\sigma}_r^2\right)^{-1}\boldsymbol{X}$$

that leads to:

$$\mathrm{var}(\boldsymbol{\beta}|\text{data}) = \sigma_\beta^2(\boldsymbol{I} - \boldsymbol{X}'(\boldsymbol{XX}' + \boldsymbol{I}\theta)^{-1}\boldsymbol{X}) \qquad (2)$$

*Estimation of the exact gametic variance:*

We aim now to predict the gametic variance of a cross, $\hat{\sigma}_{P_1\times P_2}^2$ or $\mathrm{var}\left(\left(\boldsymbol{X}_{\boldsymbol{P_1}\times\boldsymbol{P_2}}\right)^{\mathrm{RIL}(k)}\boldsymbol{\beta}|\text{data}\right)$, with $\left(\boldsymbol{X}_{\boldsymbol{P_1}\times\boldsymbol{P_2}}\right)^{\mathrm{RIL}(k)}$ the vector of genotypes for the RIL progeny of cross $P_1 \times P_2$ with $k$ generations.

We assume:

$$\mathrm{E}\left[\left(\boldsymbol{X}_{\boldsymbol{P_1}\times\boldsymbol{P_2}}\right)^{\mathrm{RIL}(k)}\boldsymbol{\beta}|\mathrm{data}\right] = \mathrm{E}\left[\left(\boldsymbol{X}_{\boldsymbol{P_1}\times\boldsymbol{P_2}}\right)^{\mathrm{RIL}(k)}\boldsymbol{\beta}|\boldsymbol{X}_{\boldsymbol{P_1}\times\boldsymbol{P_2}},\boldsymbol{X},\boldsymbol{y}\right]$$

$$= \mathrm{E}\left[\left(\boldsymbol{X}_{\boldsymbol{P_1}\times\boldsymbol{P_2}}\right)^{\mathrm{RIL}(k)}|\boldsymbol{X}_{\boldsymbol{P_1}\times\boldsymbol{P_2}},\boldsymbol{X},\boldsymbol{y}\right]\mathrm{E}[\boldsymbol{\beta}|\boldsymbol{X}_{\boldsymbol{P_1}\times\boldsymbol{P_2}},\boldsymbol{X},\boldsymbol{y}]$$

$$\mathrm{E}\left[\left(\boldsymbol{X}_{\boldsymbol{P_1}\times\boldsymbol{P_2}}\right)^{\mathrm{RIL}(k)}\boldsymbol{\beta}|\mathrm{data}\right] = 0.5\boldsymbol{X}_{\boldsymbol{P_1}\times\boldsymbol{P_2}}\widehat{\boldsymbol{\beta}} \quad (3)$$

Following (Bernardo 2014; Mohammadi et al. 2015; Lehermeier et al. 2017), an approximation of gametic variance is:

$$\hat{\sigma}^2_{P_1\times P_2} = \widehat{\boldsymbol{\beta}}'\boldsymbol{V}_{\boldsymbol{P_1}\times\boldsymbol{P_2}}\widehat{\boldsymbol{\beta}}$$

with $\boldsymbol{V}_{\boldsymbol{P_1}\times\boldsymbol{P_2}}$ the genotypic variance-covariance matrix for biparental progeny. Diagonal terms of this matrix are 0 if parents carry the same allele at *loci j* and $l$, or 0.25 if alleles are in coupling phase, that is one parent carries the two beneficial alleles while the other carries deleterious alleles, or -0.25 if the alleles are in repulsion phase), k the number of generations, and $c_{jl}$ the recombination rate between *loci j* and $l$.

To compute the exact $\mathrm{var}\left(\left(\boldsymbol{X}_{\boldsymbol{P_1}\times\boldsymbol{P_2}}\right)^{\mathrm{RIL}(k)}\boldsymbol{\beta}|\boldsymbol{X}_{\boldsymbol{P_1}\times\boldsymbol{P_2}},\boldsymbol{X},\boldsymbol{y}\right)$, we use the following property:

$\mathrm{var}(a|c) = \mathrm{var}_b(\mathrm{E}[a|b,c]) + \mathrm{E}_b[\mathrm{var}(a|b,c)]$.

With $a = \left(\boldsymbol{X}_{\boldsymbol{P_1}\times\boldsymbol{P_2}}\right)^{\mathrm{RIL}(k)}\boldsymbol{\beta}$, $b = \boldsymbol{\beta}$ and $c = \boldsymbol{X}_{\boldsymbol{P_1}\times\boldsymbol{P_2}},\boldsymbol{X},\boldsymbol{y}$, we obtain**:**

$$\mathrm{var}\left(\left(\boldsymbol{X}_{\boldsymbol{P_1}\times\boldsymbol{P_2}}\right)^{\mathrm{RIL}(k)}\boldsymbol{\beta}\middle|\boldsymbol{X}_{\boldsymbol{P_1}\times\boldsymbol{P_2}},\boldsymbol{X},\boldsymbol{y}\right) = \mathrm{var}_{\boldsymbol{\beta}}\left(\mathrm{E}\left[\left(\boldsymbol{X}_{\boldsymbol{P_1}\times\boldsymbol{P_2}}\right)^{\mathrm{RIL}(k)}\boldsymbol{\beta}|\boldsymbol{\beta},\boldsymbol{X}_{\boldsymbol{P_1}\times\boldsymbol{P_2}},\boldsymbol{X},\boldsymbol{y}\right]\right)$$

$$+\mathrm{E}_{\boldsymbol{\beta}}\left[\mathrm{var}\left(\left(\boldsymbol{X}_{\boldsymbol{P_1}\times\boldsymbol{P_2}}\right)^{\mathrm{RIL}(k)}\boldsymbol{\beta}|\boldsymbol{\beta},\boldsymbol{X}_{\boldsymbol{P_1}\times\boldsymbol{P_2}},\boldsymbol{X},\boldsymbol{y}\right)\right]$$

From equation (3) we deduce the first term:

$$\mathrm{var}_{\boldsymbol{\beta}}\left(\mathrm{E}\left[\left(\boldsymbol{X}_{\boldsymbol{P_1}\times\boldsymbol{P_2}}\right)^{\mathrm{RIL}(k)}\boldsymbol{\beta}|\mathrm{data}\right]\right) = 0.25\boldsymbol{X}_{\boldsymbol{P_1}\times\boldsymbol{P_2}}\mathrm{var}(\boldsymbol{\beta}|\mathrm{data})\boldsymbol{X}'_{\boldsymbol{P_1}\times\boldsymbol{P_2}}$$

From the following two properties, we deduce the second term:

Property 1: $\text{var}(Ma) = a'\text{var}(M)a$

Property 2: $\text{E}[a'Ma|\text{data}] = \text{E}[a|\text{data}]'M\text{E}[a|\text{data}] + \text{trace}\big(M\text{var}(a|\text{data})\big)$

$$\begin{aligned}\text{E}_{\boldsymbol{\beta}}\left[\text{var}\left(\left(\boldsymbol{X}_{\boldsymbol{P_1}\times\boldsymbol{P_2}}\right)^{\text{RIL}(k)}\boldsymbol{\beta}|\text{data}\right)\right] &= \text{E}_{\boldsymbol{\beta}}\big[\widehat{\boldsymbol{\beta}}'\boldsymbol{V}_{\boldsymbol{P_1}\times\boldsymbol{P_2}}\widehat{\boldsymbol{\beta}}\big] \\ &= \text{E}[\boldsymbol{\beta}|\text{data}]'\boldsymbol{V}_{\boldsymbol{P_1}\times\boldsymbol{P_2}}\text{E}[\boldsymbol{\beta}|\text{data}] + \text{trace}\left(\boldsymbol{V}_{\boldsymbol{P_1}\times\boldsymbol{P_2}}\text{var}(\boldsymbol{\beta}|\text{data})\right) \\ &= \widehat{\boldsymbol{\beta}}'\boldsymbol{V}_{\boldsymbol{P_1}\times\boldsymbol{P_2}}\widehat{\boldsymbol{\beta}} + \text{trace}\left(\boldsymbol{V}_{\boldsymbol{P_1}\times\boldsymbol{P_2}}\text{var}(\boldsymbol{\beta}|\text{data})\right)\end{aligned}$$

The final formula for the exact computation of $\hat{\sigma}^2_{P_1\times P_2}$ is then:

$$\hat{\sigma}^2_{P_1\times P_2} = \widehat{\boldsymbol{\beta}}'\boldsymbol{V}_{\boldsymbol{P_1}\times\boldsymbol{P_2}}\widehat{\boldsymbol{\beta}} + \text{trace}\{\boldsymbol{V}_{\boldsymbol{P_1}\times\boldsymbol{P_2}}\text{var}(\boldsymbol{\beta}|\boldsymbol{X},\boldsymbol{y})\} + 0.25\boldsymbol{X}'_{\boldsymbol{P_1}\times\boldsymbol{P_2}}\text{var}(\boldsymbol{\beta}|\boldsymbol{X},\boldsymbol{y})\boldsymbol{X}_{\boldsymbol{P_1}\times\boldsymbol{P_2}} \quad (4)$$

In our study we decompose equation (4) into 3 estimators for the variance of gametes, $\text{Vg}_1$, $\text{Vg}_2$ and $\text{Vg}_3$:

$$\left(\hat{\sigma}^2_{P_1\times P_2}\right)_{\text{Vg}_1} = \widehat{\boldsymbol{\beta}}'\boldsymbol{V}_{\boldsymbol{P_1}\times\boldsymbol{P_2}}\widehat{\boldsymbol{\beta}}$$

$\left(\hat{\sigma}^2_{P_1\times P_2}\right)_{\text{Vg}_1}$ estimator is the classical estimator where the uncertainty in the estimate of SNP effects measured by $\text{var}(\boldsymbol{\beta}|\boldsymbol{X},\boldsymbol{y})$ is neglected.

$$\left(\hat{\sigma}^2_{P_1\times P_2}\right)_{\text{Vg}_2} = \widehat{\boldsymbol{\beta}}'\boldsymbol{V}_{\boldsymbol{P_1}\times\boldsymbol{P_2}}\widehat{\boldsymbol{\beta}} + \text{trace}\{\boldsymbol{V}_{\boldsymbol{P_1}\times\boldsymbol{P_2}}\text{var}(\boldsymbol{\beta}|\boldsymbol{X},\boldsymbol{y})\}$$

$\left(\hat{\sigma}^2_{P_1\times P_2}\right)_{\text{Vg}_2}$ is an algebraic version of the Posterior Mean Variance (PMV) method described by (Lehermeier et al. 2017), i.e. the average of the variances obtained over $L$ successive samplings in the Markov chain: $\sigma^{2\,(\text{PMV})} = \frac{1}{L}\sum_{s=1}^{L}\boldsymbol{\beta}'_{(s)}\boldsymbol{V}\boldsymbol{\beta}_{(s)}$ where $\boldsymbol{\beta}_{(s)}$ is the value obtained at the $s^{\text{th}}$ sampling. For $L$ large $\sigma^{2\,(\text{PMV})} \sim \text{E}_{\boldsymbol{\beta}}\big[\text{var}\big(\boldsymbol{X}_{\boldsymbol{P_1}\times\boldsymbol{P_2}}\boldsymbol{\beta}|\text{data}\big)\big]$.

$$\left(\hat{\sigma}^2_{P_1\times P_2}\right)_{\text{Vg}_3} = \widehat{\boldsymbol{\beta}}'\boldsymbol{V}_{\boldsymbol{P_1}\times\boldsymbol{P_2}}\widehat{\boldsymbol{\beta}} + \text{trace}\{\boldsymbol{V}_{\boldsymbol{P_1}\times\boldsymbol{P_2}}\text{var}(\boldsymbol{\beta}|\boldsymbol{X},\boldsymbol{y})\} + 0.25\boldsymbol{X}'_{\boldsymbol{P_1}\times\boldsymbol{P_2}}\text{var}(\boldsymbol{\beta}|\boldsymbol{X},\boldsymbol{y})\boldsymbol{X}_{\boldsymbol{P_1}\times\boldsymbol{P_2}}$$

$\left(\hat{\sigma}^2_{P_1\times P_2}\right)_{\text{Vg}_3}$ considers the fact that the uncertainty of the estimation of marker effects is modulated for each sire by his own genomic constitution.

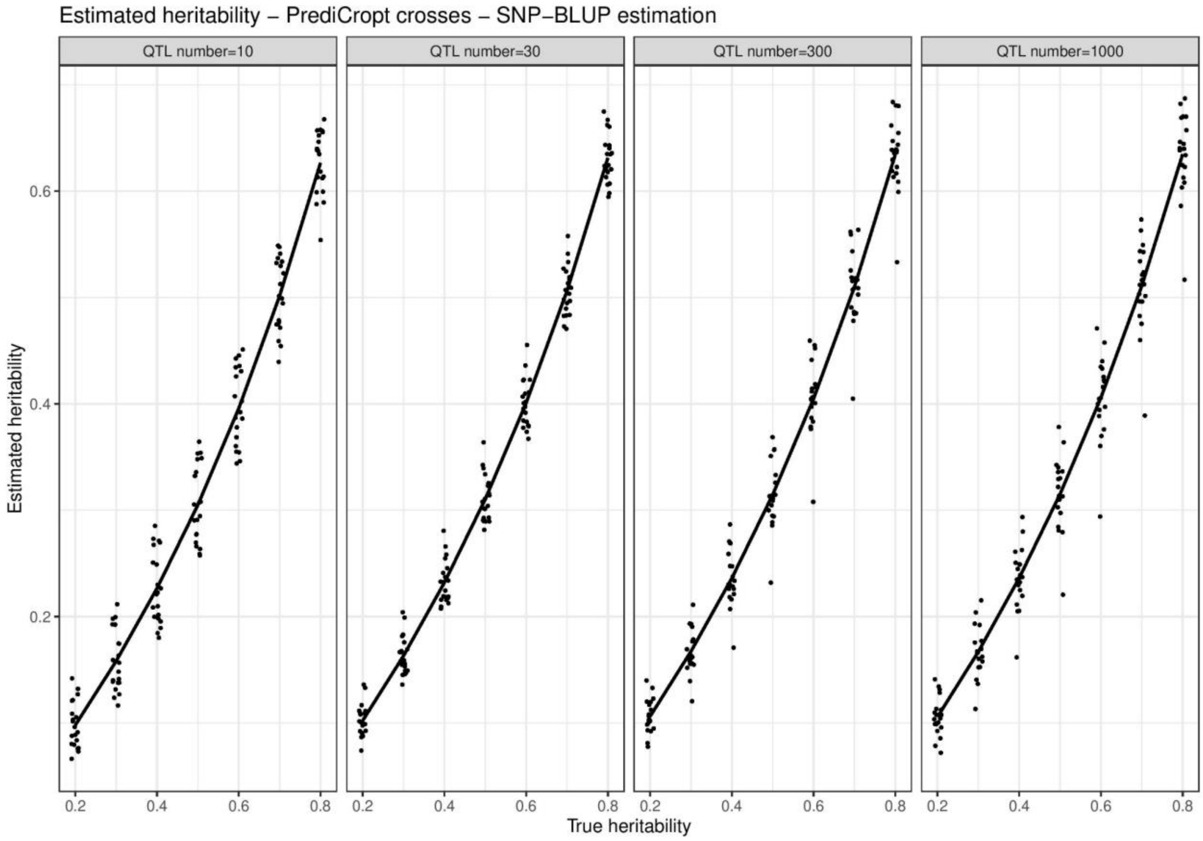


**Supplementary Figure S1. Estimated heritability as a function of true heritability in the simulation study**. *Variance components were estimated using a SNP-BLUP approach implemented in the rrBLUP R package.*

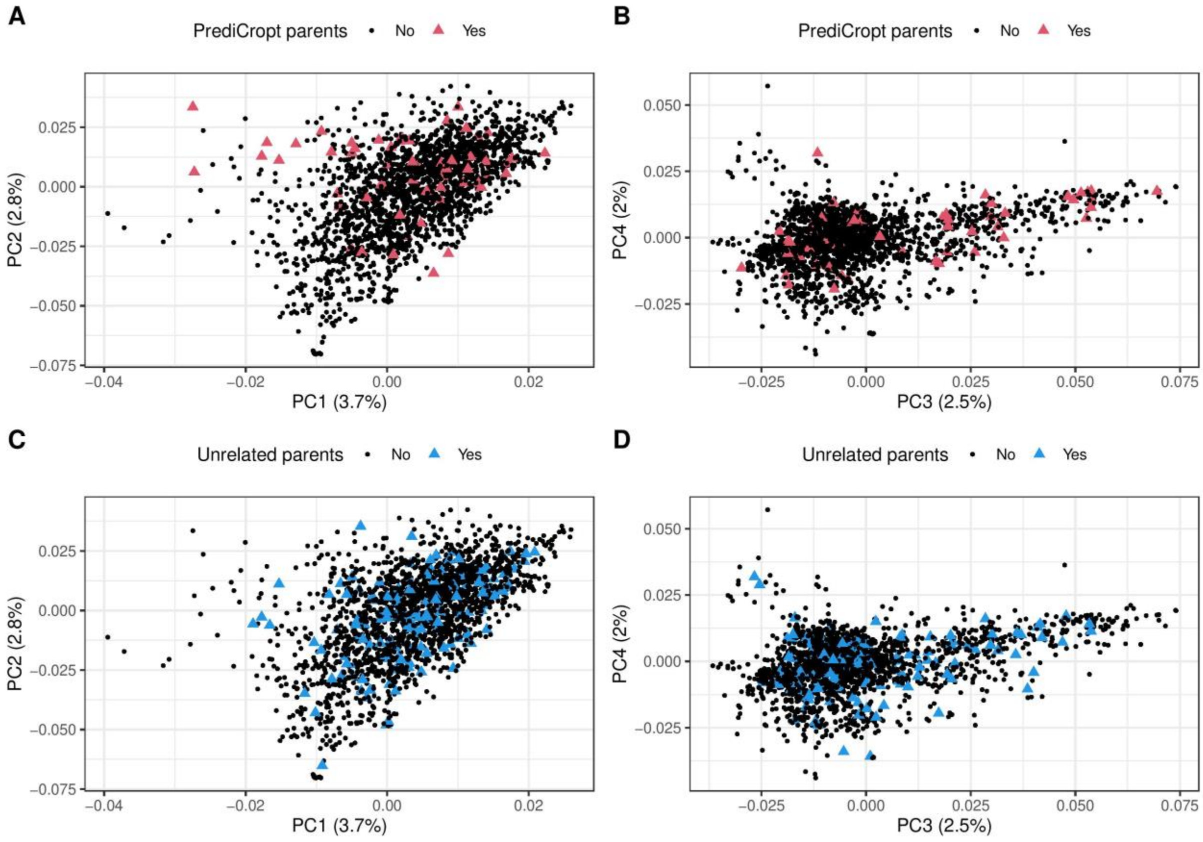


**Supplementary Figure S2. Four Principal Components (PC) decomposition for the parents of the 73 crosses of the 'PrediCropt' (A and B) and 'Unrelated' scenarii (C and D) among the phenotyped training population.**

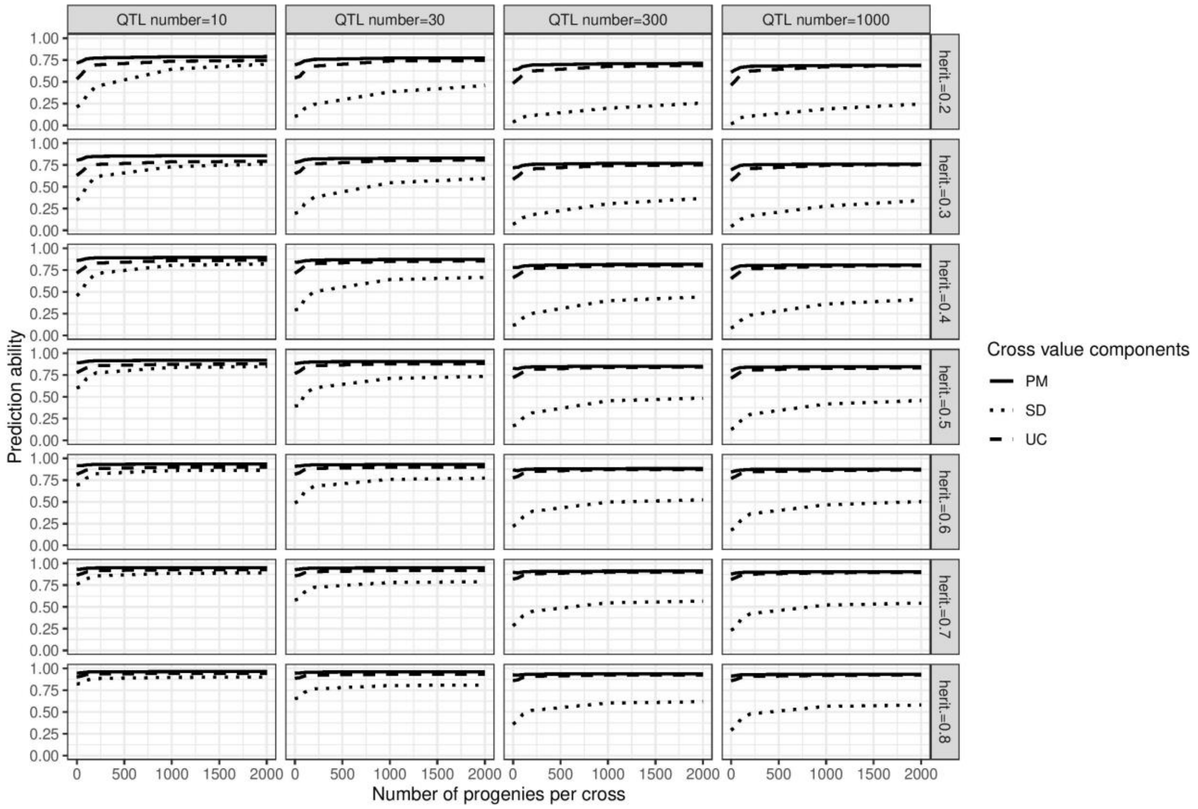


**Supplementary Figure S3. Impact of the number of progenies per cross on the prediction ability of the three cross value components, based on simulations**. *Marker effects were estimated with 'BayesA' method. Abbreviations: PM: Parental Mean, SD: Standard Deviation, UC: Usefulness Criterion.*

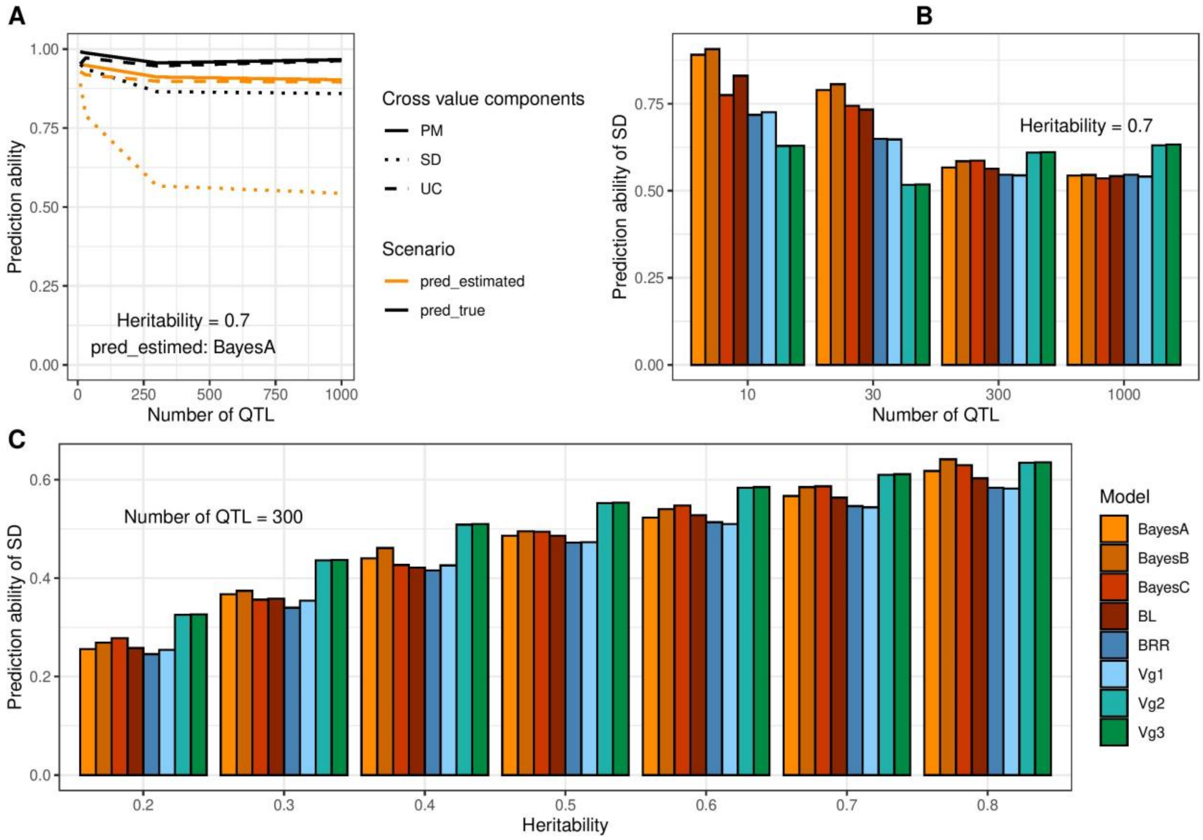


**Supplementary Figure S4. Impact of the knowledge of QTL on the prediction ability of the three cross value components (A) and of the genomic selection method on the prediction ability of SD (B and C), based on simulations**. *The number of progenies per cross was 2000 progenies per cross. In A, the heritability was 0.7 and marker effects were estimated with 'BayesA' method. The true cross value components (pred_true) were calculated with true marker effects whereas the estimated cross value components (pred_estimated) were calculated with estimated marker effects. In B, the heritability was 0.7. In C, the number of QTL was 300. Abbreviations: PM: Parental Mean, SD: Standard Deviation, UC: Usefulness Criterion.*

| Trait | Dataset | Zone | Number of environments (location*year) | Number of means | Number of lines with BLUE values | Number of lines with phenotypes and genotypes |
|---|---|---|---|---|---|---|
| Yield | GEVES | North | 548 | 19,556 | 471 | 434 |
| | | South | 297 | 8,114 | 258 | 237 |
| | INRAE-AO | North | 194 | 26,596 | 2,539 | 1,578 |
| | | South | 94 | 2,960 | 596 | 375 |
| | Both | Both | 1,031 | 57,226 | 3,241 | 2,146 |
| Grain protein content | GEVES | North | 351 | 10,807 | 449 | 416 |
| | | South | 240 | 5,616 | 257 | 236 |
| | INRAE-AO | North | 136 | 13,547 | 2,309 | 1,515 |
| | | South | 36 | 931 | 264 | 210 |
| | Both | Both | 714 | 30,901 | 2,933 | 2,062 |
| Plant height | GEVES | North | 278 | 9,314 | 469 | 432 |
| | | South | 159 | 4,231 | 258 | 237 |
| | INRAE-AO | North | 116 | 14,709 | 2,357 | 1,575 |
| | | South | 45 | 1,191 | 364 | 242 |
| | Both | Both | 551 | 29,445 | 3,008 | 2,126 |
| Heading date | GEVES | North | 448 | 15,281 | 466 | 431 |
| | | South | 215 | 5,784 | 257 | 236 |
| | INRAE-AO | North | 168 | 21,272 | 2,539 | 1,578 |
| | | South | 71 | 2,277 | 580 | 369 |
| | Both | Both | 832 | 44,614 | 3,237 | 2,145 |

**Supplementary Table S1. Description of the historical training population for the 4 analyzed traits**. *For each trial, we computed spatially adjusted means when coordinates were available and arithmetic means if not. BLUE values were then computed from these means to correct the effect of the environment (location*year).*

| North / South | 2000 | 2001 | 2002 | 2003 | 2004 | 2005 | 2006 | 2007 | 2008 | 2009 | 2010 | 2011 | 2012 | 2013 | 2014 | 2015 | 2016 | 2017 | 2018 | 2019 | 2020 | 2021 | 2022 |
|---|---|---|---|---|---|---|---|---|---|---|---|---|---|---|---|---|---|---|---|---|---|---|---|
| **2000** | 38<br>18 | 26 (0.72) | 5 | 5 | 6 | 6 | 6 | 4 | 3 | 2 | 3 | 2 | 2 | 2 | 2 | 1 | | | | | | | |
| **2001** | 10 (0.82) | 50<br>16 | 28 (0.79) | 7 | 6 | 6 | 6 | 4 | 3 | 1 | 2 | 2 | 2 | 2 | 2 | 1 | | | | | | | |
| **2002** | 3 | 9 (0.96) | 48<br>15 | 26 (0.82) | 7 | 5 | 5 | 8 | 7 | 4 | 3 | 3 | 3 | 3 | 3 | 2 | | | 1 | | | | |
| **2003** | 3 | 4 | 10 (0.90) | 44<br>15 | 24 (0.87) | 5 | 5 | 8 | 8 | 5 | 3 | 3 | 3 | 3 | 3 | 2 | | | 1 | | | | |
| **2004** | 2 | 3 | 3 | 8 (0.76) | 51<br>17 | 27 (0.86) | 6 | 5 | 4 | 3 | 4 | 3 | 3 | 3 | 3 | 2 | | | 1 | | | | |
| **2005** | 2 | 3 | 3 | 3 | 12 (0.82) | 50<br>19 | 29 (0.69) | 7 | 5 | 4 | 6 | 6 | 6 | 5 | 4 | 3 | | | 1 | | | | |
| **2006** | 2 | 3 | 3 | 3 | 5 | 11 (0.80) | 47<br>20 | 25 (0.73) | 5 | 4 | 6 | 6 | 6 | 5 | 5 | 4 | 2 | 1 | 2 | | | | |
| **2007** | 3 | 4 | 3 | 3 | 3 | 3 | 12 (0.85) | 52<br>25 | 30 (0.67) | 5 | 4 | 3 | 3 | 3 | 4 | 3 | 2 | 1 | 2 | | | | |
| **2008** | 2 | 3 | 2 | 3 | 4 | 3 | 3 | 15 (0.80) | 50<br>26 | 24 (0.67) | 3 | 2 | 2 | 2 | 2 | 1 | | | 1 | | | | |
| **2009** | 2 | 3 | 2 | 3 | 4 | 3 | 3 | 3 | 14 (0.66) | 52<br>27 | 30 (0.78) | 4 | 3 | 3 | 2 | 2 | 1 | 1 | 2 | 1 | | 1 | 1 |
| **2010** | 3 | 4 | 3 | 3 | 3 | 3 | 3 | 4 | 3 | 16 (0.77) | 54<br>34 | 28 (0.91) | 5 | 5 | 5 | 6 | 3 | 3 | 4 | 2 | 1 | 2 | 2 |
| **2011** | 3 | 4 | 3 | 3 | 3 | 3 | 3 | 4 | 3 | 4 | 22 (0.87) | 55<br>38 | 29 (0.62) | 5 | 5 | 5 | 2 | 3 | 3 | 1 | 1 | 1 | 1 |
| **2012** | 3 | 4 | 3 | 3 | 3 | 3 | 3 | 5 | 4 | 3 | 6 | 22 (0.77) | 54<br>39 | 27 (0.83) | 4 | 3 | 1 | 2 | 2 | 1 | 1 | 1 | |
| **2013** | 3 | 4 | 3 | 3 | 3 | 3 | 3 | 5 | 4 | 3 | 5 | 6 | 23 (0.89) | 52<br>32 | 28 (0.87) | 4 | 1 | 1 | 3 | 1 | 1 | 1 | |
| **2014** | 2 | 3 | 3 | 3 | 3 | 3 | 3 | 4 | 3 | 3 | 4 | 5 | 6 | 15 (0.85) | 46<br>33 | 20 (0.86) | 2 | 1 | 3 | 1 | 2 | 2 | 1 |
| **2015** | 1 | 2 | 2 | 2 | 2 | 2 | 2 | 3 | 2 | 1 | 3 | 4 | 4 | 4 | 21 (0.89) | 42<br>38 | 24 (0.50) | 3 | 4 | 3 | 3 | 4 | 3 |
| **2016** | | | | | | | | 2 | 2 | 1 | 2 | 3 | 2 | 2 | 3 | 19 (0.73) | 49<br>33 | 27 (0.25) | 6 | 4 | 5 | 6 | 4 |
| **2017** | | | | | | | | 2 | 2 | 1 | 2 | 2 | 1 | 1 | 2 | 2 | 16 (0.69) | 44<br>27 | 21 (0.80) | 3 | 4 | 6 | 4 |
| **2018** | | | | | | | | | | | 1 | 1 | | | 1 | 4 | 3 | 12 (0.91) | 38<br>30 | 18 (0.49) | 2 | 4 | 4 |
| **2019** | | | | | | | | | | | 1 | 1 | | | 3 | 6 | 3 | 1 | 19 (0.71) | 34<br>29 | 18 (0.82) | 6 | 6 |
| **2020** | | | | | | | | | | | 1 | 1 | | | 3 | 5 | 3 | 2 | 3 | 13 (0.77) | 41<br>32 | 26 (0.87) | 6 |
| **2021** | | | | | | | | | | | 1 | 1 | | | 2 | 4 | 3 | 2 | 2 | 3 | 22 (0.78) | 43<br>33 | 23 (0.89) |
| **2022** | | | | | | | | | | | 1 | 1 | | | 3 | 5 | 3 | 2 | 3 | 4 | 5 | 16 (0.90) | 25<br>18 |

**Supplementary Table S2. Number of lines for yield in GEVES dataset per year and in common between years, and correlation between years**. *Correlations are provided in parenthesis for consecutive years only. North dataset is above the diagonal and South, below. An empty cell means 0.*

| North<br>South | 2000 | 2001 | 2002 | 2003 | 2004 | 2005 | 2006 | 2007 | 2008 | 2009 | 2010 | 2011 | 2012 | 2013 | 2014 | 2015 | 2016 | 2017 | 2018 | 2019 | 2020 | 2021 | 2022 |
|---|---|---|---|---|---|---|---|---|---|---|---|---|---|---|---|---|---|---|---|---|---|---|---|
| **2000** | 144<br>24 | 46<br>(0.61) | 24 | 15 | 7 | 5 | 4 | 2 | 2 | 2 | 2 | 2 | 2 | 2 | 2 | 2 | 1 | 1 | 2 | 1 | 1 | 1 | 1 |
| **2001** | 8<br>(0.09) | 166<br>27 | 59<br>(0.74) | 29 | 16 | 13 | 5 | 2 | 2 | 3 | 3 | 2 | 2 | 2 | 2 | 1 | 1 | | 1 | | | | |
| **2002** | 5 | 13<br>(0.90) | 184<br>24 | 59<br>(0.54) | 27 | 18 | 7 | 4 | 4 | 6 | 5 | 3 | 3 | 3 | 3 | 2 | 1 | 1 | 2 | 1 | 1 | 1 | 1 |
| **2003** | 3 | 6 | 10<br>(0.82) | 185<br>22 | 57<br>(0.70) | 24 | 11 | 5 | 5 | 6 | 6 | 4 | 4 | 4 | 4 | 3 | 1 | 1 | 2 | 1 | 1 | 1 | 1 |
| **2004** | 3 | 4 | 5 | 8<br>(0.94) | 181<br>26 | 52<br>(0.78) | 21 | 11 | 8 | 7 | 7 | 5 | 5 | 5 | 5 | 4 | 1 | 1 | 2 | 1 | 1 | 1 | 1 |
| **2005** | 3 | 6 | 6 | 6 | 9<br>(0.88) | 164<br>19 | 56<br>(0.86) | 24 | 11 | 8 | 8 | 5 | 5 | 5 | 5 | 4 | 1 | 1 | 2 | 1 | 1 | 1 | 1 |
| **2006** | 3 | 5 | 4 | 4 | 7 | 11<br>(0.81) | 161<br>28 | 59<br>(0.27) | 24 | 11 | 9 | 5 | 6 | 5 | 5 | 5 | 2 | 1 | 2 | 1 | 1 | 1 | 1 |
| **2007** | 2 | 3 | 3 | 3 | 3 | 4 | 7<br>(0.83) | 178<br>26 | 73<br>(0.52) | 29 | 14 | 7 | 8 | 6 | 6 | 5 | 2 | 1 | 2 | 1 | 1 | 1 | 1 |
| **2008** | 2 | 3 | 3 | 3 | 3 | 3 | 4 | 10<br>(0.80) | 174<br>47 | 57<br>(0.72) | 29 | 13 | 11 | 8 | 8 | 7 | 4 | 1 | 2 | 1 | 1 | 1 | 1 |
| **2009** | 2 | 2 | 2 | 2 | 3 | 3 | 4 | 8 | 14<br>(0.62) | 172<br>64 | 55<br>(0.71) | 23 | 13 | 8 | 8 | 6 | 5 | 2 | 3 | 2 | 1 | 1 | 1 |
| **2010** | 1 | 1 | 1 | 1 | 1 | 1 | 2 | 3 | 5 | 13<br>(-0.07) | 174<br>59 | 67<br>(0.65) | 32 | 17 | 13 | 11 | 6 | 4 | 3 | 2 | 1 | 1 | 1 |
| **2011** | 1 | 1 | 1 | 1 | 1 | 1 | 1 | 2 | 2 | 3 | 14<br>(0.10) | 178<br>42 | 61<br>(0.43) | 31 | 21 | 16 | 8 | 4 | 3 | 2 | 1 | 1 | 1 |
| **2012** | 1 | 1 | 1 | 1 | 1 | 1 | 1 | 2 | 2 | 2 | 4 | 10<br>(0.59) | 181<br>30 | 66<br>(0.54) | 39 | 17 | 9 | 5 | 3 | 2 | 1 | 1 | 1 |
| **2013** | 1 | 1 | 1 | 1 | 1 | 1 | 2 | 3 | 3 | 3 | 3 | 4 | 14<br>(0.33) | 176<br>35 | 84<br>(0.74) | 38 | 20 | 9 | 7 | 4 | 1 | 2 | 1 |
| **2014** | 1 | 2 | 2 | 2 | 2 | 2 | 3 | 3 | 3 | 2 | 2 | 2 | 6 | 16<br>(0.86) | 176<br>52 | 77<br>(0.81) | 37 | 18 | 8 | 6 | 1 | 2 | 1 |
| **2015** | 1 | 2 | 2 | 2 | 2 | 2 | 2 | 2 | 2 | 1 | 1 | 2 | 3 | 8 | 19<br>(0.78) | 183<br>45 | 79<br>(00.52) | 37 | 15 | 7 | 2 | 3 | 2 |
| **2016** | | | | | | | | | | | | 1 | 2 | 5 | 12 | 22<br>(0.83) | 157<br>49 | 67<br>(0.41) | 24 | 12 | 4 | 2 | 1 |
| **2017** | | | | | | | | | | | | | 1 | 1 | 6 | 12 | 23<br>(0.34) | 178<br>65 | 67<br>(0.69) | 23 | 9 | 5 | 4 |
| **2018** | | | | | | | | | | | | | | | 2 | 7 | 10 | 25<br>(0.64) | 154<br>46 | 40<br>(0.50) | 10 | 5 | 4 |
| **2019** | 1 | 1 | 1 | 1 | 1 | 1 | 1 | 1 | 1 | 1 | 1 | 1 | 1 | 1 | 1 | 8 | 8 | 10 | 14<br>(0.69) | 180<br>60 | 45<br>(0.51) | 18 | 12 |
| **2020** | 1 | 1 | 1 | 1 | 1 | 1 | 1 | 1 | 1 | 1 | 1 | 1 | 1 | 1 | 1 | 6 | 6 | 8 | 9 | 25<br>(0.23) | 172<br>52 | 56<br>(0.55) | 20 |
| **2021** | 1 | 1 | 1 | 1 | 1 | 1 | 1 | 1 | 1 | 1 | 1 | 1 | 1 | 1 | 1 | 2 | 2 | 3 | 3 | 13 | 26<br>(0.57) | 144<br>48 | 36<br>(0.67) |
| **2022** | 1 | 1 | 1 | 1 | 1 | 1 | 1 | 1 | 1 | 1 | 1 | 1 | 1 | 1 | 1 | 2 | 1 | 1 | 1 | 10 | 15 | 27<br>(0.46) | 120<br>58 |

**Supplementary Table S3. Number of lines for yield in INRAE-AO dataset per year and in common between years, and correlation between years**.

*Correlations are provided in parenthesis for consecutive years only. North dataset is above the diagonal and South, below. An empty cell means 0.*

| North / South | 2000 | 2001 | 2002 | 2003 | 2004 | 2005 | 2006 | 2007 | 2008 | 2009 | 2010 | 2011 | 2012 | 2013 | 2014 | 2015 | 2016 | 2017 | 2018 | 2019 | 2020 | 2021 | 2022 |
|---|---|---|---|---|---|---|---|---|---|---|---|---|---|---|---|---|---|---|---|---|---|---|---|
| **2000** | 36 / 18 | 24 (0.55) | 5 | 5 | 6 | 5 | 5 | 3 | 2 | 2 | 3 | 2 | 2 | 2 | 2 | 1 | | | | | | | |
| **2001** | 10 (0.85) | 49 / 16 | 28 (0.38) | 6 | 5 | 4 | 4 | 2 | 1 | 1 | 2 | 2 | 2 | 2 | 2 | 1 | | | | | | | |
| **2002** | 3 | 9 (0.72) | 48 / 15 | 26 (0.73) | 7 | 4 | 4 | 6 | 5 | 4 | 3 | 3 | 3 | 3 | 3 | 2 | | | 1 | | | | |
| **2003** | 3 | 4 | 10 (0.94) | 43 / 15 | 24 (0.95) | 4 | 4 | 6 | 5 | 4 | 3 | 3 | 3 | 3 | 3 | 2 | | | 1 | | | | |
| **2004** | 2 | 3 | 3 | 8 (0.96) | 47 / 17 | 27 (0.92) | 6 | 4 | 3 | 3 | 4 | 3 | 3 | 3 | 3 | 2 | | | 1 | | | | |
| **2005** | 2 | 3 | 3 | 3 | 12 (0.91) | 47 / 18 | 26 (0.88) | 4 | 3 | 4 | 6 | 6 | 6 | 5 | 4 | 3 | | | 1 | | | | |
| **2006** | 2 | 3 | 3 | 3 | 5 | 11 (0.80) | 44 / 20 | 22 (0.73) | 3 | 4 | 6 | 6 | 6 | 5 | 5 | 4 | 2 | 1 | 2 | | | | |
| **2007** | 3 | 4 | 3 | 3 | 3 | 3 | 12 (0.61) | 47 / 25 | 27 (0.89) | 5 | 4 | 3 | 3 | 3 | 4 | 3 | 2 | 1 | 2 | | | | |
| **2008** | 2 | 3 | 2 | 3 | 4 | 3 | 3 | 15 (0.59) | 46 / 26 | 24 (0.72) | 3 | 2 | 2 | 2 | 2 | 1 | | | 1 | | | | |
| **2009** | 2 | 3 | 2 | 3 | 4 | 3 | 3 | 3 | 14 (0.84) | 50 / 27 | 29 (0.90) | 3 | 3 | 3 | 2 | 2 | 1 | 1 | 2 | 1 | | | |
| **2010** | 3 | 4 | 3 | 3 | 3 | 3 | 3 | 4 | 3 | 16 (0.89) | 54 / 32 | 28 (0.91) | 5 | 5 | 5 | 6 | 3 | 3 | 4 | 2 | 1 | 1 | 1 |
| **2011** | 3 | 4 | 3 | 3 | 3 | 3 | 3 | 4 | 3 | 4 | 20 (0.76) | 52 / 37 | 29 (0.82) | 5 | 5 | 5 | 2 | 3 | 3 | 1 | 1 | 1 | 1 |
| **2012** | 3 | 4 | 3 | 3 | 3 | 3 | 3 | 5 | 4 | 3 | 4 | 21 (0.88) | 51 / 39 | 27 (0.83) | 4 | 3 | 1 | 2 | 2 | 1 | 1 | 1 | |
| **2013** | 3 | 4 | 3 | 3 | 3 | 3 | 3 | 5 | 4 | 3 | 4 | 5 | 23 (0.94) | 52 / 32 | 28 (0.85) | 4 | 1 | 1 | 3 | 1 | 1 | 1 | |
| **2014** | 2 | 3 | 3 | 3 | 3 | 3 | 3 | 4 | 3 | 3 | 4 | 5 | 6 | 15 (0.94) | 44 / 33 | 20 (0.89) | 2 | 1 | 3 | 1 | 2 | 2 | 1 |
| **2015** | 1 | 2 | 2 | 2 | 2 | 2 | 2 | 3 | 2 | 1 | 3 | 4 | 4 | 4 | 21 (0.97) | 42 / 38 | 24 (0.70) | 3 | 4 | 3 | 3 | 3 | 2 |
| **2016** | | | | | | | | 2 | 2 | 1 | 2 | 3 | 2 | 2 | 3 | 19 (0.83) | 48 / 33 | 26 (0.41) | 6 | 4 | 5 | 5 | 3 |
| **2017** | | | | | | | | 2 | 2 | 1 | 2 | 2 | 1 | 1 | 2 | 2 | 16 (0.85) | 42 / 27 | 20 (0.84) | 3 | 4 | 5 | 3 |
| **2018** | | | | | | | | | | | 1 | 1 | | | 1 | 4 | 3 | 12 (0.91) | 38 / 29 | 18 (0.61) | 2 | 3 | 2 |
| **2019** | | | | | | | | | | | 1 | 1 | | | 3 | 6 | 3 | 1 | 18 (0.83) | 34 / 29 | 18 (0.83) | 5 | 4 |
| **2020** | | | | | | | | | | | 1 | 1 | | | 3 | 5 | 3 | 2 | 2 | 13 (0.93) | 38 / 32 | 25 (0.81) | 6 |
| **2021** | | | | | | | | | | | 1 | 1 | | | 2 | 4 | 3 | 2 | 2 | 3 | 22 (0.92) | 42 / 33 | 22 (0.88) |
| **2022** | | | | | | | | | | | 1 | 1 | | | 2 | 4 | 3 | 2 | 2 | 3 | 5 | 16 (0.92) | 22 / 16 |

**Supplementary Table S4. Number of lines for grain protein content in GEVES dataset per year and in common between years, and correlation between years.** *Correlations are provided in parenthesis for consecutive years only. North dataset is above the diagonal and South, below. An empty cell means 0.*

| North<br>South | 2000 | 2001 | 2002 | 2003 | 2004 | 2005 | 2006 | 2007 | 2008 | 2009 | 2010 | 2011 | 2012 | 2013 | 2014 | 2015 | 2016 | 2017 | 2018 | 2019 | 2020 | 2021 | 2022 |
|---|---|---|---|---|---|---|---|---|---|---|---|---|---|---|---|---|---|---|---|---|---|---|---|
| **2000** | 143 | | 24 | 15 | 7 | 5 | 4 | 2 | 2 | 2 | 2 | 2 | 2 | 2 | 2 | 1 | 1 | 1 | 2 | 1 | 1 | 1 | 1 |
| **2001** | | 9 | | | | | | | | | | | | | | | | | | | | | |
| **2002** | | | 64 | 27 (0.85) | 14 | 11 | 3 | 3 | 3 | 4 | 4 | 3 | 3 | 3 | 3 | 1 | 1 | 1 | 2 | 1 | 1 | 1 | 1 |
| **2003** | | | | 185 | 57 (0.80) | 24 | 11 | 5 | 5 | 6 | 6 | 4 | 4 | 4 | 4 | 2 | 1 | 1 | 2 | 1 | 1 | 1 | 1 |
| **2004** | | | | | 181 | 52 (0.62) | 21 | 11 | 8 | 7 | 7 | 5 | 5 | 5 | 5 | 2 | 1 | 1 | 2 | 1 | 1 | 1 | 1 |
| **2005** | | 2 | | | | 164<br>17 | 56 (0.85) | 24 | 11 | 8 | 8 | 5 | 5 | 5 | 5 | 2 | 1 | 1 | 2 | 1 | 1 | 1 | 1 |
| **2006** | | 2 | | | | 10 (0.86) | 161<br>28 | 59 (0.72) | 24 | 11 | 9 | 5 | 6 | 5 | 5 | 2 | 2 | 1 | 2 | 1 | 1 | 1 | 1 |
| **2007** | | | | | | | | 178 | 73 (0.72) | 29 | 14 | 7 | 8 | 6 | 6 | 2 | 2 | 1 | 2 | 1 | 1 | 1 | 1 |
| **2008** | | | | | | | | | 174 | 57 (0.80) | 29 | 13 | 11 | 8 | 8 | 2 | 4 | 1 | 2 | 1 | 1 | 1 | 1 |
| **2009** | | 2 | | | | 3 | 4 | | | 172<br>41 | 55 (0.86) | 23 | 13 | 8 | 8 | 1 | 5 | 2 | 3 | 2 | 1 | 1 | 1 |
| **2010** | | 1 | | | | 1 | 2 | | | 9 (0.83) | 174<br>23 | 67 (0.86) | 32 | 17 | 13 | 4 | 6 | 4 | 3 | 2 | 1 | 1 | 1 |
| **2011** | | 1 | | | | 1 | 1 | | | 3 | 8 (0.28) | 178<br>22 | 61 (0.72) | 31 | 21 | 7 | 8 | 4 | 3 | 2 | 1 | 1 | 1 |
| **2012** | | 1 | | | | 1 | 1 | | | 2 | 3 | 5 (-0.28) | 181<br>30 | 66 (0.73) | 39 | 8 | 9 | 5 | 3 | 2 | 1 | 1 | 1 |
| **2013** | | 1 | | | | 1 | 2 | | | 3 | 3 | 3 | 14 (0.50) | 176<br>16 | 84 (0.90) | 28 | 20 | 9 | 7 | 4 | 1 | 2 | 1 |
| **2014** | | 1 | | | | 1 | 3 | | | 2 | 2 | 2 | 6 | 8 (0.44) | 176<br>34 | 35 (0.78) | 37 | 18 | 8 | 6 | 1 | 2 | 1 |
| **2015** | | | | | | | | | | | | | | | | 40 | 17 (0.87) | 8 | 7 | 4 | 2 | 3 | 2 |
| **2016** | | | | | | | | | | | | 1 | 2 | 2 | 6 | | 157<br>26 | 67 (0.64) | 24 | 12 | 4 | 2 | 1 |
| **2017** | | | | | | | | | | | | | 1 | 1 | 2 | | 14 (0.61) | 178<br>28 | 67 (0.79) | 23 | 9 | 5 | 4 |
| **2018** | | | | | | | | | | | | | | | | | 7 | 13 (0.91) | 154<br>28 | 40 (0.91) | 10 | 5 | 4 |
| **2019** | | 1 | | | | 1 | 1 | | | 1 | 1 | 1 | 1 | 1 | 1 | | 6 | 8 | 9 (0.98) | 180<br>28 | 45 (0.86) | 18 | 12 |
| **2020** | | 1 | | | | 1 | 1 | | | 1 | 1 | 1 | 1 | 1 | 1 | | 4 | 6 | 6 | 11 (0.98) | 172<br>28 | 56 (0.83) | 20 |
| **2021** | | | | | | | | | | | | | | | | | 1 | 2 | 3 | 4 | 16 (0.84) | 144<br>22 | 36 (0.84) |
| **2022** | | | | | | | | | | | | | | | | | 1 | 1 | 1 | 2 | 9 | 13 (0.86) | 120<br>32 |

**Supplementary Table S5. Number of lines for grain protein content in INRAE-AO dataset per year and in common between years, and correlation between years**. *Correlations are provided in parenthesis for consecutive years only. North dataset is above the diagonal and South, below. An empty cell means 0.*

| North / South | 2000 | 2001 | 2002 | 2003 | 2004 | 2005 | 2006 | 2007 | 2008 | 2009 | 2010 | 2011 | 2012 | 2013 | 2014 | 2015 | 2016 | 2017 | 2018 | 2019 | 2020 | 2021 | 2022 |
|---|---|---|---|---|---|---|---|---|---|---|---|---|---|---|---|---|---|---|---|---|---|---|---|
| **2000** | 38 / 18 | 26 (0.94) | 5 | 5 | 6 | 5 | 6 | 4 | 2 | 2 | 3 | 2 | 2 | 2 | 2 | 1 | | | | | | | |
| **2001** | 10 (0.93) | 50 / 16 | 28 (0.95) | 7 | 6 | 4 | 6 | 3 | 1 | 1 | 2 | 2 | 2 | 2 | 2 | 1 | | | | | | | |
| **2002** | 3 | 9 (0.96) | 48 / 15 | 26 (0.94) | 7 | 4 | 5 | 7 | 5 | 4 | 3 | 3 | 3 | 3 | 3 | 2 | | | 1 | | | | |
| **2003** | 3 | 4 | 10 (0.93) | 44 / 15 | 24 (0.97) | 4 | 5 | 7 | 6 | 5 | 3 | 3 | 3 | 3 | 3 | 2 | | | 1 | | | | |
| **2004** | 2 | 3 | 3 | 8 (0.89) | 51 / 17 | 27 (0.97) | 6 | 5 | 3 | 3 | 4 | 3 | 3 | 3 | 3 | 2 | | | 1 | | | | |
| **2005** | 2 | 3 | 3 | 3 | 12 (0.96) | 47 / 19 | 26 (0.98) | 5 | 3 | 4 | 6 | 6 | 6 | 5 | 4 | 3 | | | 1 | | | | |
| **2006** | 2 | 3 | 3 | 3 | 5 | 11 (0.92) | 47 / 20 | 25 (0.95) | 3 | 4 | 6 | 6 | 6 | 5 | 5 | 4 | 2 | 1 | 2 | | | | |
| **2007** | 3 | 4 | 3 | 3 | 3 | 3 | 12 (0.89) | 32 / 25 | 28 (0.94) | 5 | 4 | 3 | 3 | 3 | 3 | 2 | | | 1 | | | | |
| **2008** | 2 | 3 | 2 | 3 | 4 | 3 | 3 | 15 (0.88) | 47 / 26 | 24 (0.94) | 3 | 2 | 2 | 2 | 2 | 1 | | | 1 | | | | |
| **2009** | 2 | 3 | 2 | 3 | 4 | 3 | 3 | 3 | 14 (0.85) | 52 / 27 | 30 (0.97) | 4 | 3 | 3 | 2 | 2 | 1 | 1 | 2 | 1 | | 1 | |
| **2010** | 3 | 4 | 3 | 3 | 3 | 3 | 3 | 4 | 3 | 16 (0.97) | 54 / 34 | 28 (0.95) | 5 | 5 | 5 | 6 | 3 | 3 | 4 | 2 | 1 | 2 | 1 |
| **2011** | 3 | 4 | 3 | 3 | 3 | 3 | 3 | 4 | 3 | 4 | 22 (0.94) | 55 / 38 | 29 (0.87) | 5 | 5 | 5 | 2 | 3 | 3 | 1 | 1 | 1 | 1 |
| **2012** | 3 | 4 | 3 | 3 | 3 | 3 | 3 | 5 | 4 | 3 | 6 | 22 (0.92) | 54 / 39 | 27 (0.95) | 4 | 3 | 1 | 2 | 2 | 1 | 1 | 1 | |
| **2013** | 3 | 4 | 3 | 3 | 3 | 3 | 3 | 5 | 4 | 3 | 5 | 6 | 23 (0.85) | 52 / 23 | 28 (0.94) | 4 | 1 | 1 | 3 | 1 | 1 | 1 | |
| **2014** | 2 | 3 | 3 | 3 | 3 | 3 | 3 | 4 | 3 | 3 | 4 | 5 | 6 | 6 (0.89) | 46 / 33 | 20 (0.93) | 2 | 1 | 3 | 1 | 2 | 2 | 1 |
| **2015** | 1 | 2 | 2 | 2 | 2 | 2 | 2 | 3 | 2 | 1 | 3 | 4 | 4 | 4 | 21 (0.94) | 42 / 38 | 24 (0.93) | 3 | 4 | 3 | 3 | 4 | 2 |
| **2016** | | | | | | | | 2 | 2 | 1 | 2 | 3 | 2 | 2 | 3 | 19 (0.85) | 49 / 33 | 27 (0.82) | 6 | 4 | 5 | 6 | 3 |
| **2017** | | | | | | | | 2 | 2 | 1 | 2 | 2 | 1 | 1 | 2 | 2 | 16 (0.91) | 44 / 27 | 21 (0.95) | 3 | 4 | 6 | 3 |
| **2018** | | | | | | | | | | | 1 | 1 | | | 1 | 4 | 3 | 12 (0.96) | 38 / 30 | 18 (0.94) | 2 | 4 | 2 |
| **2019** | | | | | | | | | | | 1 | 1 | | | 3 | 6 | 3 | 1 | 19 (0.91) | 34 / 29 | 18 (0.92) | 6 | 4 |
| **2020** | | | | | | | | | | | 1 | 1 | | | 3 | 5 | 3 | 2 | 3 | 13 (0.78) | 41 / 32 | 26 (0.96) | 6 |
| **2021** | | | | | | | | | | | 1 | 1 | | | 2 | 4 | 3 | 2 | 2 | 3 | 22 (0.89) | 43 / 33 | 22 (0.42) |
| **2022** | | | | | | | | | | | 1 | 1 | | | 2 | 4 | 3 | 2 | 2 | 3 | 5 | 16 (0.77) | 22 / 16 |

**Supplementary Table S6. Number of lines for plant height in GEVES dataset per year and in common between years, and correlation between years**.

*Correlations are provided in parenthesis for consecutive years only. North dataset is above the diagonal and South, below. An empty cell means 0.*

| North<br>South | 2000 | 2001 | 2002 | 2003 | 2004 | 2005 | 2006 | 2007 | 2008 | 2009 | 2010 | 2011 | 2012 | 2013 | 2014 | 2015 | 2016 | 2017 | 2018 | 2019 | 2020 | 2021 | 2022 |
|---|---|---|---|---|---|---|---|---|---|---|---|---|---|---|---|---|---|---|---|---|---|---|---|
| **2000** | 144 | 46 (0.72) | 24 | 15 | | 5 | 4 | 2 | 2 | 2 | 2 | 2 | 2 | 2 | 2 | 2 | 1 | 1 | 2 | 1 | 1 | 1 | 1 |
| **2001** | | 166<br>26 | 58 (0.37) | 29 | | 13 | 5 | 2 | 2 | 3 | 3 | 2 | 2 | 2 | 2 | 1 | 1 | | 1 | | | | |
| **2002** | | 13 (0.93) | 64<br>24 | 27 (0.80) | | 11 | 3 | 3 | 3 | 4 | 4 | 3 | 3 | 3 | 3 | 2 | 1 | 1 | 2 | 1 | 1 | 1 | 1 |
| **2003** | | 6 | 10 (0.88) | 185<br>21 | | 24 | 11 | 5 | 5 | 6 | 6 | 4 | 4 | 4 | 4 | 3 | 1 | 1 | 2 | 1 | 1 | 1 | 1 |
| **2004** | | 4 | 5 | 8 (0.92) | 26 | | | | | | | | | | | | | | | | | | |
| **2005** | | 4 | 5 | 5 | 9 (0.93) | 164<br>18 | 56 (0.92) | 24 | 11 | 8 | 8 | 5 | 5 | 5 | 5 | 4 | 1 | 1 | 2 | 1 | 1 | 1 | 1 |
| **2006** | | 4 | 4 | 4 | 7 | 11 (0.78) | 161<br>28 | 59 (0.86) | 24 | 11 | 9 | 5 | 6 | 5 | 5 | 5 | 2 | 1 | 2 | 1 | 1 | 1 | 1 |
| **2007** | | 3 | 3 | 3 | 3 | 4 | 7 (0.67) | 178<br>26 | 73 (0.90) | 29 | 14 | 7 | 8 | 6 | 6 | 5 | 2 | 1 | 2 | 1 | 1 | 1 | 1 |
| **2008** | | 3 | 3 | 3 | 3 | 3 | 4 | 10 (0.67) | 174<br>47 | 57 (0.87) | 29 | 13 | 11 | 8 | 8 | 7 | 4 | 1 | 2 | 1 | 1 | 1 | 1 |
| **2009** | | 2 | 2 | 2 | 3 | 3 | 4 | 8 | 14 (0.74) | 172<br>41 | 55 (0.91) | 23 | 13 | 8 | 8 | 6 | 5 | 2 | 3 | 2 | 1 | 1 | 1 |
| **2010** | | 1 | 1 | 1 | 1 | 1 | 2 | 3 | 5 | 9 (0.86) | 174<br>23 | 67 (0.84) | 32 | 17 | 13 | 11 | 6 | 4 | 3 | 2 | 1 | 1 | 1 |
| **2011** | | 1 | 1 | 1 | 1 | 1 | 1 | 2 | 2 | 3 | 8 (0.89) | 178<br>22 | 61 (0.88) | 31 | 21 | 16 | 8 | 4 | 3 | 2 | 1 | 1 | 1 |
| **2012** | | 1 | 1 | 1 | 1 | 1 | 1 | 2 | 2 | 2 | 3 | 5 (0.97) | 181<br>30 | 66 (0.82) | 39 | 17 | 9 | 5 | 3 | 2 | 1 | 1 | 1 |
| **2013** | | 1 | 1 | 1 | 1 | 1 | 2 | 3 | 3 | 3 | 3 | 3 | 14 (0.62) | 176<br>16 | 84 (0.87) | 38 | 20 | 9 | 7 | 4 | 1 | 2 | 1 |
| **2014** | | 2 | 2 | 2 | 2 | 2 | 3 | 3 | 3 | 2 | 2 | 2 | 6 | 8 (0.87) | 176<br>34 | 77 (0.93) | 37 | 18 | 8 | 6 | 1 | 2 | 1 |
| **2015** | | 2 | 2 | 2 | 2 | 2 | 2 | 2 | 2 | 1 | 1 | 2 | 3 | 4 | 11 (0.93) | 183<br>25 | 79 (0.87) | 37 | 15 | 7 | 2 | 3 | 2 |
| **2016** | | | | | | | | | | | | 1 | 2 | 2 | 6 | 13 (0.84) | 157<br>26 | 67 (0.88) | 24 | 12 | 4 | 2 | 1 |
| **2017** | | | | | | | | | | | | | 1 | 1 | 2 | 8 | 14 (0.63) | 178<br>28 | 67 (0.93) | 23 | 9 | 5 | 4 |
| **2018** | | | | | | | | | | | | | | | | 5 | 7 | 13 (0.92) | 154<br>28 | 40 (0.90) | 10 | 5 | 4 |
| **2019** | | 1 | 1 | 1 | 1 | 1 | 1 | 1 | 1 | 1 | 1 | 1 | 1 | 1 | 1 | 4 | 6 | 8 | 9 (0.93) | 180<br>28 | 45 (0.84) | 18 | 12 |
| **2020** | | 1 | 1 | 1 | 1 | 1 | 1 | 1 | 1 | 1 | 1 | 1 | 1 | 1 | 1 | 4 | 4 | 6 | 6 | 11 (0.59) | 172<br>28 | 56 (0.87) | 20 |
| **2021** | | | | | | | | | | | | | | | | 1 | 1 | 2 | 3 | 4 | 16 (0.69) | 144<br>28 | 36 (0.88) |
| **2022** | | | | | | | | | | | | | | | | 1 | 1 | 1 | 1 | 2 | 9 | 17 (0.66) | 120<br>32 |

**Supplementary Table S7. Number of lines for plant height in INRAE-AO dataset per year and in common between years, and correlation between years**.

*Correlations are provided in parenthesis for consecutive years only. North dataset is above the diagonal and South, below. An empty cell means 0.*

| North / South | 2000 | 2001 | 2002 | 2003 | 2004 | 2005 | 2006 | 2007 | 2008 | 2009 | 2010 | 2011 | 2012 | 2013 | 2014 | 2015 | 2016 | 2017 | 2018 | 2019 | 2020 | 2021 | 2022 |
|---|---|---|---|---|---|---|---|---|---|---|---|---|---|---|---|---|---|---|---|---|---|---|---|
| **2000** | 38 / 18 | 26 (0.97) | 5 | 5 | 6 | 6 | 6 | 4 | 2 | 2 | 3 | 2 | 2 | 2 | 2 | 1 | | | | | | | |
| **2001** | 10 (0.97) | 50 / 16 | 28 (0.98) | 7 | 6 | 6 | 6 | 4 | 1 | 1 | 2 | 2 | 2 | 2 | 2 | 1 | | | | | | | |
| **2002** | 3 | 9 (0.94) | 48 / 15 | 26 (0.96) | 7 | 5 | 5 | 8 | 5 | 4 | 3 | 3 | 3 | 3 | 3 | 2 | | | 1 | | | | |
| **2003** | 3 | 4 | 10 (0.96) | 44 / 15 | 24 (0.97) | 5 | 5 | 8 | 6 | 5 | 3 | 3 | 3 | 3 | 3 | 2 | | | 1 | | | | |
| **2004** | 2 | 3 | 3 | 8 (0.92) | 36 / 17 | 15 (0.99) | 6 | 5 | 3 | 3 | 4 | 3 | 3 | 3 | 3 | 2 | | | 1 | | | | |
| **2005** | | | | | | 49 | 28 (0.80) | 6 | 3 | 4 | 6 | 6 | 6 | 5 | 4 | 3 | | | 1 | | | | |
| **2006** | 2 | 3 | 3 | 3 | 5 | | 47 / 20 | 25 (0.90) | 3 | 4 | 6 | 6 | 6 | 5 | 5 | 4 | 2 | 1 | 2 | | | | |
| **2007** | 3 | 4 | 3 | 3 | 3 | | 3 (1.00) | 52 / 16 | 28 (0.97) | 5 | 4 | 3 | 3 | 3 | 4 | 3 | 2 | 1 | 2 | | | | |
| **2008** | 2 | 3 | 2 | 3 | 4 | | 3 | 15 (0.81) | 47 / 26 | 24 (0.97) | 3 | 2 | 2 | 2 | 2 | 1 | | | 1 | | | | |
| **2009** | 2 | 3 | 2 | 3 | 4 | | 3 | 3 | 14 (0.95) | 52 / 27 | 30 (0.98) | 4 | 3 | 3 | 2 | 2 | 1 | 1 | 2 | 1 | | 1 | |
| **2010** | 3 | 4 | 3 | 3 | 3 | | 3 | 4 | 3 | 16 (0.90) | 54 / 34 | 28 (0.97) | 5 | 5 | 5 | 6 | 3 | 3 | 4 | 2 | 1 | 2 | 1 |
| **2011** | 3 | 4 | 3 | 3 | 3 | | 3 | 4 | 3 | 4 | 22 (0.95) | 55 / 38 | 29 (0.97) | 5 | 5 | 5 | 2 | 3 | 3 | 1 | 1 | 1 | 1 |
| **2012** | 3 | 4 | 3 | 3 | 3 | | 3 | 5 | 4 | 3 | 6 | 22 (0.88) | 54 / 39 | 27 (0.96) | 4 | 3 | 1 | 2 | 2 | 1 | 1 | 1 | |
| **2013** | 3 | 4 | 3 | 3 | 3 | | 3 | 5 | 4 | 3 | 5 | 6 | 23 (0.94) | 52 / 23 | 28 (0.97) | 4 | 1 | 1 | 3 | 1 | 1 | 1 | |
| **2014** | 2 | 3 | 3 | 3 | 3 | | 3 | 4 | 3 | 3 | 4 | 5 | 6 | 6 (0.99) | 46 / 33 | 20 (0.97) | 2 | 1 | 3 | 1 | 2 | 2 | 1 |
| **2015** | 1 | 2 | 2 | 2 | 2 | | 2 | 3 | 2 | 1 | 3 | 4 | 4 | 4 | 21 (0.95) | 42 / 38 | 24 (0.97) | 3 | 4 | 3 | 3 | 4 | 2 |
| **2016** | | | | | | | | 2 | 2 | 1 | 2 | 3 | 2 | 2 | 3 | 19 (0.93) | 49 / 33 | 27 (0.89) | 6 | 4 | 5 | 6 | 3 |
| **2017** | | | | | | | | 2 | 2 | 1 | 2 | 2 | 1 | 1 | 2 | 2 | 16 (0.55) | 44 / 27 | 21 (0.96) | 3 | 4 | 6 | 3 |
| **2018** | | | | | | | | | | | 1 | 1 | | | 1 | 4 | 3 | 12 (0.97) | 38 / 30 | 18 (0.99) | 2 | 4 | 2 |
| **2019** | | | | | | | | | | | 1 | 1 | | | 3 | 6 | 3 | 1 | 19 (0.92) | 34 / 29 | 18 (0.97) | 6 | 4 |
| **2020** | | | | | | | | | | | 1 | 1 | | | 3 | 5 | 3 | 2 | 3 | 13 (0.91) | 41 / 32 | 26 (0.97) | 6 |
| **2021** | | | | | | | | | | | 1 | 1 | | | 2 | 4 | 3 | 2 | 2 | 3 | 22 (0.93) | 43 / 33 | 22 (0.96) |
| **2022** | | | | | | | | | | | 1 | 1 | | | 2 | 4 | 3 | 2 | 2 | 3 | 5 | 16 (0.89) | 22 / 16 |

**Supplementary Table S8. Number of lines for heading date in GEVES dataset per year and in common between years, and correlation between years**.

*Correlations are provided in parenthesis for consecutive years only. North dataset is above the diagonal and South, below. An empty cell means 0.*

| North<br>South | 2000 | 2001 | 2002 | 2003 | 2004 | 2005 | 2006 | 2007 | 2008 | 2009 | 2010 | 2011 | 2012 | 2013 | 2014 | 2015 | 2016 | 2017 | 2018 | 2019 | 2020 | 2021 | 2022 |
|---|---|---|---|---|---|---|---|---|---|---|---|---|---|---|---|---|---|---|---|---|---|---|---|
| **2000** | 144 | 46 (0.89) | 24 | 15 | 7 | 5 | 4 | 2 | 2 | 2 | 2 | 2 | 2 | 2 | 2 | 2 | 1 | 1 | 2 | 1 | 1 | 1 | 1 |
| **2001** | | 166<br>26 | 59 (0.93) | 29 | 16 | 13 | 5 | 2 | 2 | 3 | 3 | 2 | 2 | 2 | 2 | 1 | 1 | | 1 | | | | |
| **2002** | | 13 (0.97) | 184<br>24 | 59 (0.88) | 27 | 18 | 7 | 4 | 4 | 6 | 5 | 3 | 3 | 3 | 3 | 2 | 1 | 1 | 2 | 1 | 1 | 1 | 1 |
| **2003** | | 6 | 10 (0.95) | 185<br>21 | 57 (0.86) | 24 | 11 | 5 | 5 | 6 | 6 | 4 | 4 | 4 | 4 | 3 | 1 | 1 | 2 | 1 | 1 | 1 | 1 |
| **2004** | | 4 | 5 | 8 (0.88) | 181<br>26 | 52 (0.76) | 21 | 11 | 8 | 7 | 7 | 5 | 5 | 5 | 5 | 4 | 1 | 1 | 2 | 1 | 1 | 1 | 1 |
| **2005** | | 4 | 5 | 5 | 9 (0.88) | 164<br>18 | 56 (0.92) | 24 | 11 | 8 | 8 | 5 | 5 | 5 | 5 | 4 | 1 | 1 | 2 | 1 | 1 | 1 | 1 |
| **2006** | | 4 | 4 | 4 | 7 | 11 (0.95) | 161<br>28 | 59 (0.80) | 24 | 11 | 9 | 5 | 6 | 5 | 5 | 5 | 2 | 1 | 2 | 1 | 1 | 1 | 1 |
| **2007** | | 3 | 3 | 3 | 3 | 4 | 7 (0.94) | 178<br>26 | 73 (0.93) | 29 | 14 | 7 | 8 | 6 | 6 | 5 | 2 | 1 | 2 | 1 | 1 | 1 | 1 |
| **2008** | | 3 | 3 | 3 | 3 | 3 | 4 | 10 (0.97) | 174<br>47 | 57 (0.80) | 29 | 13 | 11 | 8 | 8 | 7 | 4 | 1 | 2 | 1 | 1 | 1 | 1 |
| **2009** | | 2 | 2 | 2 | 3 | 3 | 4 | 8 | 14 (0.89) | 172<br>64 | 55 (0.95) | 23 | 13 | 8 | 8 | 6 | 5 | 2 | 3 | 2 | 1 | 1 | 1 |
| **2010** | | 1 | 1 | 1 | 1 | 1 | 2 | 3 | 5 | 13 (0.47) | 174<br>59 | 67 (0.87) | 32 | 17 | 13 | 11 | 6 | 4 | 3 | 2 | 1 | 1 | 1 |
| **2011** | | 1 | 1 | 1 | 1 | 1 | 1 | 2 | 2 | 3 | 14 (0.90) | 178<br>42 | 61 (0.94) | 31 | 21 | 16 | 8 | 4 | 3 | 2 | 1 | 1 | 1 |
| **2012** | | 1 | 1 | 1 | 1 | 1 | 1 | 2 | 2 | 2 | 4 | 10 (0.77) | 181<br>30 | 66 (0.92) | 39 | 17 | 9 | 5 | 3 | 2 | 1 | 1 | 1 |
| **2013** | | 1 | 1 | 1 | 1 | 1 | 2 | 3 | 3 | 3 | 3 | 4 | 14 (0.96) | 176<br>35 | 84 (0.88) | 38 | 20 | 9 | 7 | 4 | 1 | 2 | 1 |
| **2014** | | 2 | 2 | 2 | 2 | 2 | 3 | 3 | 3 | 2 | 2 | 2 | 6 | 16 (0.81) | 176<br>52 | 77 (0.94) | 37 | 18 | 8 | 6 | 1 | 2 | 1 |
| **2015** | | 2 | 2 | 2 | 2 | 2 | 2 | 2 | 2 | 1 | 1 | 2 | 3 | 8 | 19 (0.89) | 183<br>45 | 79 (0.95) | 37 | 15 | 7 | 2 | 3 | 2 |
| **2016** | | | | | | | | | | | | 1 | 2 | 5 | 12 | 22 (0.79) | 157<br>49 | 67 (0.94) | 24 | 12 | 4 | 2 | 1 |
| **2017** | | | | | | | | | | | | | 1 | 1 | 6 | 12 | 23 (0.91) | 178<br>65 | 67 (0.97) | 23 | 9 | 5 | 4 |
| **2018** | | | | | | | | | | | | | | | 2 | 7 | 10 | 25 (0.97) | 154<br>46 | 40 (0.95) | 10 | 5 | 4 |
| **2019** | | 1 | 1 | 1 | 1 | 1 | 1 | 1 | 1 | 1 | 1 | 1 | 1 | 1 | 1 | 8 | 8 | 10 | 14 (0.70) | 180<br>60 | 45 (0.94) | 18 | 12 |
| **2020** | | 1 | 1 | 1 | 1 | 1 | 1 | 1 | 1 | 1 | 1 | 1 | 1 | 1 | 1 | 6 | 6 | 8 | 9 | 25 (0.76) | 172<br>52 | 56 (0.95) | 20 |
| **2021** | | 1 | 1 | 1 | 1 | 1 | 1 | 1 | 1 | 1 | 1 | 1 | 1 | 1 | 1 | 2 | 2 | 3 | 3 | 13 | 26 (0.80) | 144<br>48 | 36 (0.96) |
| **2022** | | 1 | 1 | 1 | 1 | 1 | 1 | 1 | 1 | 1 | 1 | 1 | 1 | 1 | 1 | 2 | 1 | 1 | 1 | 10 | 15 | 27 (0.94) | 120<br>58 |

**Supplementary Table S9. Number of lines for heading date in INRAE-AO dataset per year and in common between years, and correlation between years.** *Correlations are provided in parenthesis for consecutive years only. North dataset is above the diagonal and South, below. An empty cell means 0.*

| Organism | Type of data | Location or group | Raw 2020 | Raw 2021 | Raw 2022 | Clean Yield | Clean Grain protein content | Clean Plant height | Clean Heading date |
|---|---|---|---|---|---|---|---|---|---|
| FD | Plots | CAP | 457 | 327 | 0 | 782 | 784 | 784 | 784 |
| | | HOU | 459 | 364 | 301 | 1,124 | 1,123 | 1,124 | 1,124 |
| | | Parents | 210 | 178 | 94 | 481 | 481 | 482 | 482 |
| | | Progenies | 706 | 513 | 207 | 1,425 | 1,426 | 1,426 | 1,426 |
| | | Total | 916 | 691 | 301 | 1,906 | 1,907 | 1,908 | 1,908 |
| | Lines | Parents | 14 | 18 | 28 | 29 | 29 | 29 | 29 |
| | | Progenies | 666 | 473 | 207 | 1,139 | 1,139 | 1,139 | 1,139 |
| | | Total | 680 | 491 | 235 | 1,168 | 1,168 | 1,168 | 1,168 |
| INRAE | Plots | AUZ | 419 | 0 | 820 | 1,236 | 408 | 1,239 | 1,239 |
| | | CLE | 0 | 800 | 804 | 1,591 | 789 | 1,604 | 1,604 |
| | | LUS | 420 | 834 | 1,003 | 2,257 | 1,246 | 2,257 | 2,255 |
| | | MON | 660 | 802 | 1,008 | 2,270 | 1,283 | 2,294 | 2,294 |
| | | Parents | 93 | 508 | 549 | 1,141 | 591 | 1,142 | 1,142 |
| | | Progenies | 1,406 | 1,928 | 3,086 | 6,213 | 3,135 | 6,252 | 6,250 |
| | | Total | 1,499 | 2,436 | 3,635 | 7,354 | 3,726 | 7,394 | 7,392 |
| | Lines | Parents | 2 | 52 | 74 | 74 | 52 | 74 | 74 |
| | | Progenies | 818 | 1,735 | 2,698 | 4,426 | 2,383 | 4,452 | 4,451 |
| | | Total | 820 | 1,787 | 2,772 | 4,500 | 2,435 | 4,526 | 4,525 |
| Both | Adjusted phenotypes | Total | | | | 7,867 | 4,638 | 7,907 | 7,905 |
| | BLUE | Total | | | | 5,658 | 3,596 | 5,684 | 5,683 |

**Supplementary Table S10. Description of the experimental data for the 4 analyzed traits**.

*Abbreviations for the protocol's locations are the following: CAP: Cappelle, HOU: Houville, AUZ: Auzeville, CLE: Clermont-Ferrand, LUS: Lusignan, MON: Mons. Clean data is after discussion with partners concerning the quality of the observations. BLUE values were computed from spatially adjusted clean observations to correct the effect of the environment (location*year).*

| P1 | P2 | Yield | | | | Grain protein content | | | | Plant height | | | | Heading date | | | |
|---|---|---|---|---|---|---|---|---|---|---|---|---|---|---|---|---|---|
| | | Nb progenies | Mean | SD | Observed UC | Nb progenies | Mean | SD | Observed UC | Nb progenies | Mean | SD | UC | Nb progenies | Mean | SD | Observed UC |
| **2** | 78 | 22 | 93.7 | 4.7 | 103.1 | 18 | 11.3 | 0.8 | 12.9 | 22 | 87.8 | 5.2 | 98.1 | 22 | 132.9 | 1.7 | 135.1 |
| **3** | 51 | 26 | 85.1 | 9.5 | 99.7 | 21 | 12.4 | 0.9 | 13.8 | 26 | 88.9 | 21.7 | 126.8 | 25 | 138.6 | 4.9 | 144.0 |
| **5** | 107 | 69 | 86.9 | 5.8 | 98.7 | 52 | 11.8 | 0.9 | 13.6 | 73 | 91.0 | 11.5 | 112.1 | 73 | 136.8 | 4.0 | 144.4 |
| **7** | 18 | 37 | 88.1 | 8.7 | 100.9 | 37 | 11.7 | 0.6 | 12.8 | 37 | 89.0 | 6.5 | 101.9 | 37 | 135.2 | 4.6 | 142.3 |
| **7** | 41 | 82 | 92.4 | 8.3 | 107.2 | 82 | 11.7 | 0.6 | 12.7 | 82 | 99.4 | 8.0 | 115.6 | 82 | 136.7 | 3.5 | 143.0 |
| **7** | 50 | 24 | 84.8 | 7.8 | 97.3 | 24 | 11.4 | 0.5 | 12.3 | 24 | 91.2 | 6.4 | 102.4 | 24 | 140.1 | 3.1 | 146.7 |
| **7** | 55 | 26 | 82.1 | 9.2 | 97.0 | 26 | 12.1 | 0.7 | 13.7 | 26 | 89.9 | 17.2 | 115.4 | 26 | 137.5 | 6.9 | 151.6 |
| **7** | 79 | 44 | 84.3 | 8.8 | 99.1 | 34 | 11.8 | 1.1 | 15.2 | 44 | 85.0 | 8.3 | 103.1 | 44 | 135.8 | 4.1 | 143.6 |
| **7** | 84 | 34 | 82.8 | 8.6 | 96.1 | 34 | 11.9 | 0.7 | 13.2 | 34 | 83.2 | 16.2 | 111.5 | 34 | 137.6 | 5.7 | 150.6 |
| **7** | 85 | 60 | 85.5 | 6.0 | 94.8 | 60 | 11.4 | 0.4 | 12.2 | 60 | 91.5 | 5.2 | 102.1 | 60 | 136.1 | 3.4 | 143.1 |
| **7** | 88 | 10 | 89.1 | 10.8 | 99.5 | 12 | 11.5 | 0.5 | 12.2 | 13 | 95.4 | 14.7 | 122.9 | 13 | 135.1 | 4.5 | 143.7 |
| **7** | 100 | 36 | 89.0 | 6.3 | 100.4 | 36 | 11.4 | 0.7 | 12.6 | 36 | 94.1 | 5.9 | 104.1 | 36 | 138.7 | 7.3 | 146.4 |
| **7** | 106 | 19 | 84.6 | 8.5 | 99.1 | 19 | 11.9 | 0.7 | 13.7 | 19 | 92.2 | 18.4 | 123.8 | 19 | 137.1 | 5.2 | 146.4 |
| **8** | 37 | 68 | 84.4 | 6.5 | 95.0 | 53 | 12.0 | 1.0 | 14.4 | 68 | 90.6 | 15.5 | 114.9 | 68 | 139.0 | 3.5 | 146.5 |
| **8** | 58 | 59 | 84.8 | 5.5 | 95.0 | 48 | 12.4 | 0.9 | 14.5 | 59 | 88.0 | 6.5 | 100.0 | 59 | 132.1 | 2.9 | 136.7 |
| **8** | 105 | 62 | 89.0 | 5.9 | 98.7 | 47 | 11.7 | 0.8 | 12.9 | 62 | 95.2 | 10.9 | 115.5 | 62 | 135.4 | 3.7 | 143.5 |
| **12** | 2 | 49 | 86.9 | 6.3 | 98.1 | 38 | 11.4 | 0.6 | 12.5 | 49 | 86.2 | 5.6 | 96.8 | 49 | 132.9 | 3.3 | 139.7 |
| **12** | 7 | 31 | 91.3 | 7.9 | 105.5 | 22 | 11.6 | 0.8 | 12.9 | 31 | 87.3 | 10.3 | 114.0 | 31 | 133.5 | 4.6 | 142.6 |
| **12** | 88 | 60 | 89.6 | 8.2 | 104.6 | 46 | 11.3 | 0.8 | 13.1 | 60 | 86.0 | 7.5 | 102.2 | 60 | 132.8 | 2.7 | 138.3 |
| **13** | 6 | 40 | 87.4 | 6.3 | 98.9 | 0 | | | | 40 | 99.9 | 7.8 | 113.3 | 40 | 141.0 | 3.3 | 146.5 |
| **15** | 8 | 84 | 78.8 | 10.0 | 92.7 | 83 | 12.8 | 0.7 | 14.1 | 84 | 98.2 | 10.7 | 117.3 | 84 | 134.7 | 5.2 | 145.7 |
| **16** | 20 | 43 | 79.1 | 6.8 | 90.1 | 43 | 11.9 | 0.5 | 13.1 | 44 | 102.5 | 11.1 | 124.6 | 44 | 140.9 | 3.1 | 145.6 |
| **18** | 35 | 60 | 84.6 | 6.9 | 96.7 | 0 | | | | 60 | 93.9 | 10.3 | 115.9 | 60 | 138.2 | 5.1 | 145.9 |
| **18** | 106 | 60 | 87.4 | 7.4 | 100.6 | 46 | 11.3 | 0.8 | 13.1 | 60 | 86.9 | 15.6 | 116.2 | 60 | 134.7 | 3.0 | 142.2 |
| **19** | 82 | 64 | 82.9 | 8.4 | 94.7 | 0 | | | | 64 | 91.2 | 7.1 | 105.1 | 64 | 133.7 | 3.2 | 142.1 |
| **21** | 50 | 73 | 84.5 | 5.5 | 93.2 | 53 | 12.1 | 0.9 | 13.8 | 73 | 94.6 | 14.3 | 115.9 | 73 | 139.2 | 2.0 | 142.8 |
| **25** | 50 | 68 | 85.8 | 5.0 | 94.2 | 54 | 11.7 | 1.0 | 13.8 | 68 | 96.8 | 11.1 | 117.8 | 68 | 135.2 | 3.9 | 141.3 |
| **26** | 24 | 59 | 83.8 | 8.9 | 97.7 | 0 | | | | 60 | 97.2 | 11.9 | 116.3 | 60 | 141.4 | 3.8 | 146.4 |
| **28** | 81 | 65 | 83.1 | 7.9 | 95.8 | 64 | 12.9 | 0.9 | 14.7 | 66 | 86.8 | 10.9 | 105.8 | 66 | 133.5 | 3.5 | 139.2 |
| **30** | 62 | 76 | 83.4 | 5.1 | 93.4 | 76 | 12.5 | 0.5 | 13.6 | 76 | 95.3 | 5.5 | 105.0 | 76 | 133.2 | 2.9 | 139.6 |
| **33** | 4 | 35 | 82.1 | 7.5 | 95.7 | 0 | | | | 35 | 101.3 | 8.9 | 115.9 | 35 | 145.3 | 4.0 | 151.1 |
| **36** | 106 | 48 | 90.1 | 4.7 | 98.7 | 39 | 11.1 | 0.8 | 13.1 | 48 | 89.2 | 6.7 | 100.9 | 48 | 134.7 | 2.8 | 141.4 |
| **37** | 7 | 63 | 79.6 | 9.0 | 94.8 | 0 | | | | 63 | 96.6 | 16.1 | 119.6 | 63 | 143.5 | 2.1 | 147.4 |
| **39** | 46 | 53 | 81.1 | 8.0 | 94.9 | 0 | | | | 53 | 98.8 | 15.3 | 128.6 | 53 | 139.4 | 4.4 | 147.9 |
| **40** | 88 | 8 | 89.5 | 4.9 | 96.2 | 5 | 11.4 | 0.9 | 12.6 | 8 | 87.5 | 5.8 | 92.9 | 8 | 132.1 | 1.5 | 135.3 |
| **42** | 7 | 72 | 90.7 | 6.5 | 104.4 | 0 | | | | 72 | 90.2 | 5.8 | 100.5 | 72 | 138.3 | 3.9 | 145.5 |
| **42** | 50 | 63 | 89.8 | 5.2 | 100.7 | 63 | 11.6 | 0.4 | 12.6 | 63 | 87.0 | 7.2 | 103.3 | 63 | 137.1 | 3.0 | 143.1 |
| **42** | 55 | 43 | 85.3 | 7.6 | 102.4 | 0 | | | | 44 | 94.6 | 7.7 | 109.8 | 44 | 135.5 | 4.5 | 144.5 |
| **42** | 68 | 67 | 85.9 | 8.2 | 100.0 | 0 | | | | 67 | 96.2 | 13.6 | 117.7 | 67 | 136.2 | 4.5 | 147.1 |
| **42** | 102 | 75 | 88.6 | 6.6 | 101.1 | 0 | | | | 75 | 89.6 | 5.4 | 98.8 | 75 | 138.7 | 3.5 | 145.9 |
| **45** | 48 | 84 | 92.9 | 6.4 | 105.3 | 84 | 11.3 | 0.4 | 12.1 | 84 | 92.4 | 5.1 | 102.4 | 84 | 138.6 | 2.8 | 143.3 |
| **47** | 29 | 84 | 89.5 | 5.6 | 99.4 | 84 | 11.6 | 0.4 | 12.5 | 84 | 91.2 | 5.2 | 101.4 | 84 | 134.6 | 2.3 | 139.7 |
| **50** | 92 | 60 | 91.1 | 5.6 | 100.2 | 60 | 11.1 | 0.4 | 11.9 | 60 | 89.7 | 5.7 | 100.2 | 60 | 139.4 | 1.5 | 142.3 |
| **52** | 10 | 39 | 87.9 | 9.4 | 101.6 | 0 | | | | 40 | 94.4 | 6.7 | 105.1 | 40 | 143.1 | 3.8 | 148.5 |
| **52** | 73 | 60 | 87.7 | 11.1 | 109.4 | 0 | | | | 60 | 93.5 | 6.7 | 107.8 | 60 | 141.5 | 3.2 | 146.5 |
| **52** | 108 | 59 | 91.0 | 6.5 | 104.6 | 0 | | | | 60 | 91.5 | 6.3 | 105.1 | 60 | 140.8 | 3.8 | 147.4 |
| **54** | 10 | 49 | 80.7 | 8.0 | 95.8 | 0 | | | | 50 | 98.0 | 10.1 | 113.9 | 50 | 144.3 | 2.4 | 148.3 |
| **54** | 52 | 60 | 86.4 | 9.4 | 105.0 | 0 | | | | 60 | 97.7 | 9.8 | 112.9 | 60 | 141.1 | 4.2 | 148.1 |
| **54** | 73 | 50 | 83.9 | 7.8 | 96.7 | 0 | | | | 50 | 99.6 | 7.8 | 111.9 | 50 | 141.3 | 2.1 | 145.0 |
| **54** | 108 | 50 | 81.9 | 7.5 | 94.7 | 0 | | | | 50 | 99.1 | 11.2 | 119.9 | 50 | 143.5 | 2.6 | 148.1 |
| **55** | 56 | 59 | 90.6 | 8.1 | 107.6 | 0 | | | | 59 | 86.8 | 6.8 | 101.1 | 59 | 129.6 | 2.9 | 136.4 |
| **55** | 75 | 13 | 89.7 | 6.9 | 98.1 | 8 | 11.2 | 0.6 | 11.9 | 13 | 84.8 | 3.4 | 92.2 | 13 | 132.3 | 1.6 | 135.1 |
| **55** | 101 | 61 | 82.2 | 7.6 | 95.3 | 0 | | | | 63 | 97.1 | 15.3 | 123.3 | 63 | 136.6 | 6.1 | 146.3 |
| **57** | 78 | 25 | 93.6 | 6.1 | 104.7 | 21 | 10.9 | 0.7 | 12.3 | 25 | 83.8 | 5.3 | 93.3 | 25 | 132.4 | 2.1 | 134.9 |
| **58** | 102 | 49 | 84.4 | 7.6 | 96.1 | 0 | | | | 49 | 90.3 | 7.8 | 106.9 | 49 | 132.4 | 4.3 | 140.9 |
| **59** | 38 | 60 | 80.1 | 8.4 | 92.8 | 0 | | | | 60 | 87.0 | 6.1 | 98.4 | 60 | 137.7 | 4.5 | 144.9 |
| **60** | 27 | 59 | 85.7 | 7.4 | 98.4 | 0 | | | | 59 | 99.0 | 11.1 | 117.4 | 59 | 136.8 | 3.6 | 143.9 |
| **60** | 84 | 41 | 91.4 | 5.8 | 101.1 | 32 | 11.5 | 0.8 | 12.7 | 43 | 93.0 | 6.5 | 106.2 | 43 | 135.0 | 2.2 | 137.9 |
| **62** | 1 | 72 | 85.8 | 6.4 | 97.0 | 53 | 12.3 | 0.9 | 14.4 | 73 | 95.8 | 13.8 | 116.7 | 73 | 137.1 | 4.0 | 144.5 |
| **62** | 76 | 56 | 89.0 | 7.8 | 105.2 | 0 | | | | 56 | 89.6 | 12.7 | 109.6 | 56 | 135.5 | 2.6 | 141.6 |
| **62** | 107 | 68 | 86.6 | 5.7 | 95.9 | 50 | 11.8 | 0.8 | 13.3 | 68 | 96.5 | 14.2 | 118.0 | 68 | 136.6 | 4.1 | 142.2 |
| **63** | 78 | 22 | 93.9 | 4.3 | 101.0 | 19 | 10.6 | 0.6 | 11.6 | 22 | 86.2 | 5.1 | 96.7 | 22 | 132.8 | 1.8 | 135.7 |
| **64** | 34 | 75 | 83.5 | 7.8 | 96.0 | 58 | 12.3 | 0.9 | 14.1 | 75 | 87.1 | 6.2 | 98.1 | 75 | 139.0 | 3.9 | 146.1 |
| **64** | 67 | 64 | 86.1 | 4.7 | 95.2 | 47 | 12.4 | 0.9 | 14.0 | 64 | 93.0 | 12.0 | 112.5 | 64 | 135.6 | 1.8 | 139.2 |
| **65** | 7 | 57 | 86.1 | 11.8 | 102.8 | 57 | 11.6 | 0.7 | 13.3 | 57 | 91.4 | 7.8 | 105.1 | 57 | 140.6 | 3.9 | 148.8 |
| **65** | 46 | 60 | 84.0 | 7.7 | 100.9 | 0 | | | | 60 | 89.5 | 5.3 | 98.9 | 60 | 141.7 | 3.8 | 147.9 |
| **65** | 101 | 50 | 86.0 | 10.0 | 97.0 | 49 | 11.6 | 0.6 | 12.8 | 50 | 92.4 | 9.2 | 108.2 | 50 | 140.0 | 2.1 | 143.5 |
| **66** | 44 | 85 | 87.4 | 11.1 | 101.0 | 85 | 11.6 | 0.6 | 13.1 | 85 | 88.6 | 4.9 | 97.5 | 85 | 137.6 | 3.5 | 143.1 |
| **69** | 23 | 65 | 83.6 | 6.1 | 96.0 | 51 | 11.7 | 1.1 | 13.9 | 65 | 94.9 | 9.1 | 111.9 | 65 | 136.5 | 4.3 | 143.0 |
| **70** | 96 | 81 | 84.0 | 8.0 | 96.6 | 81 | 12.6 | 0.7 | 14.2 | 81 | 93.2 | 6.8 | 105.0 | 81 | 136.4 | 3.8 | 143.7 |
| **71** | 32 | 51 | 84.7 | 4.3 | 94.2 | 39 | 11.8 | 1.1 | 13.5 | 51 | 91.2 | 4.0 | 99.6 | 51 | 133.5 | 3.4 | 140.3 |
| **71** | 69 | 68 | 86.7 | 8.3 | 97.8 | 52 | 11.1 | 1.0 | 12.6 | 68 | 92.1 | 12.0 | 115.0 | 68 | 131.5 | 6.4 | 141.6 |
| **72** | 17 | 26 | 80.6 | 7.9 | 93.0 | 18 | 12.3 | 0.9 | 14.0 | 26 | 88.5 | 17.7 | 119.6 | 26 | 140.8 | 1.6 | 142.6 |
| **72** | 94 | 18 | 83.7 | 3.6 | 92.5 | 12 | 11.9 | 0.6 | 13.0 | 18 | 97.2 | 8.0 | 120.1 | 18 | 142.3 | 2.3 | 146.5 |
| **73** | 10 | 49 | 84.4 | 11.3 | 103.4 | 0 | | | | 49 | 94.7 | 5.3 | 106.1 | 49 | 143.3 | 2.9 | 147.9 |
| **74** | 21 | 61 | 90.7 | 4.9 | 98.7 | 47 | 11.7 | 0.7 | 13.5 | 61 | 88.2 | 5.3 | 97.7 | 61 | 137.4 | 3.2 | 143.4 |
| **74** | 64 | 67 | 86.1 | 7.0 | 96.0 | 48 | 12.3 | 0.8 | 14.1 | 67 | 89.8 | 5.9 | 101.0 | 67 | 137.3 | 2.3 | 142.0 |
| **74** | 71 | 68 | 88.3 | 6.0 | 98.1 | 49 | 11.7 | 0.7 | 13.3 | 70 | 90.0 | 5.0 | 99.2 | 70 | 134.8 | 4.7 | 143.3 |
| **74** | 89 | 64 | 90.0 | 4.7 | 99.9 | 51 | 11.4 | 0.7 | 13.0 | 65 | 92.4 | 5.3 | 102.7 | 65 | 132.8 | 1.6 | 135.5 |
| **80** | 9 | 86 | 84.4 | 7.4 | 98.1 | 86 | 11.8 | 0.4 | 12.8 | 86 | 90.3 | 6.0 | 102.0 | 86 | 139.7 | 2.0 | 142.8 |

| | | | | | | | | | | | | | | | | | |
|---|---|---|---|---|---|---|---|---|---|---|---|---|---|---|---|---|---|
| **81** | 53 | 122 | 86.5 | 5.6 | 97.1 | 122 | 12.8 | 0.7 | 14.1 | 122 | 91.3 | 9.1 | 107.5 | 122 | 135.1 | 2.3 | 140.0 |
| **84** | 91 | 60 | 89.9 | 6.8 | 99.5 | 60 | 11.4 | 0.4 | 12.2 | 60 | 92.4 | 4.2 | 101.6 | 60 | 135.1 | 1.3 | 137.7 |
| **85** | 31 | 59 | 84.1 | 7.8 | 96.9 | 0 | | | | 60 | 93.2 | 12.1 | 116.7 | 60 | 134.6 | 4.1 | 144.0 |
| **85** | 49 | 81 | 86.4 | 6.4 | 96.6 | 81 | 11.6 | 0.4 | 12.4 | 81 | 90.3 | 4.3 | 97.9 | 81 | 137.9 | 2.9 | 143.6 |
| **86** | 14 | 60 | 88.1 | 6.8 | 97.9 | 60 | 11.5 | 0.5 | 12.5 | 60 | 87.8 | 4.8 | 97.0 | 60 | 136.2 | 2.0 | 139.4 |
| **87** | 11 | 60 | 88.2 | 6.3 | 99.1 | 60 | 11.8 | 0.4 | 12.5 | 60 | 88.5 | 4.5 | 97.5 | 60 | 136.2 | 1.3 | 138.6 |
| **88** | 2 | 8 | 92.5 | 8.2 | 102.2 | 6 | 11.3 | 0.7 | 12.7 | 8 | 86.3 | 7.0 | 98.2 | 8 | 134.0 | 1.9 | 135.9 |
| **90** | 5 | 50 | 89.2 | 5.0 | 100.0 | 40 | 11.5 | 0.5 | 12.8 | 50 | 90.4 | 5.8 | 102.1 | 50 | 135.9 | 2.0 | 138.9 |
| **90** | 36 | 31 | 88.9 | 4.9 | 96.5 | 31 | 11.9 | 0.6 | 12.9 | 31 | 88.0 | 4.2 | 95.1 | 31 | 133.7 | 2.5 | 137.1 |
| **95** | 102 | 59 | 80.6 | 7.8 | 92.8 | 0 | | | | 59 | 90.6 | 12.7 | 109.9 | 59 | 142.1 | 2.8 | 146.5 |
| **97** | 93 | 53 | 89.0 | 5.6 | 99.5 | 53 | 11.7 | 0.4 | 12.4 | 53 | 91.1 | 5.0 | 100.8 | 53 | 133.3 | 1.6 | 136.0 |
| **98** | 25 | 70 | 87.0 | 5.2 | 95.5 | 53 | 11.6 | 0.8 | 13.1 | 70 | 85.7 | 6.2 | 96.9 | 70 | 136.7 | 3.5 | 142.9 |
| **99** | 61 | 60 | 88.2 | 6.2 | 100.9 | 60 | 11.2 | 0.5 | 12.2 | 60 | 86.9 | 5.8 | 96.9 | 60 | 139.5 | 1.3 | 142.4 |
| **101** | 22 | 37 | 84.8 | 6.3 | 96.8 | 37 | 12.5 | 0.7 | 13.8 | 37 | 92.5 | 13.7 | 112.6 | 37 | 139.9 | 3.6 | 145.0 |
| **101** | 60 | 17 | 83.3 | 6.9 | 92.5 | 16 | 12.2 | 0.9 | 14.5 | 17 | 89.0 | 14.2 | 118.8 | 17 | 138.7 | 4.4 | 144.9 |
| **102** | 43 | 60 | 88.1 | 5.6 | 96.3 | 60 | 11.4 | 0.5 | 12.3 | 60 | 89.9 | 5.5 | 100.4 | 60 | 139.8 | 1.5 | 142.3 |
| **103** | 7 | 23 | 85.6 | 10.5 | 99.0 | 20 | 11.8 | 1.2 | 15.0 | 23 | 95.0 | 15.8 | 126.0 | 23 | 137.5 | 5.2 | 145.1 |
| **103** | 83 | 54 | 92.5 | 6.9 | 105.6 | 40 | 11.2 | 0.8 | 12.4 | 54 | 87.8 | 6.3 | 97.9 | 54 | 134.2 | 1.9 | 138.4 |
| **104** | 77 | 83 | 91.9 | 4.9 | 99.6 | 83 | 11.3 | 0.4 | 12.0 | 83 | 91.3 | 4.8 | 98.8 | 83 | 135.2 | 2.2 | 139.3 |
| **105** | 76 | 66 | 88.0 | 5.6 | 99.1 | 52 | 11.7 | 1.0 | 13.3 | 69 | 91.8 | 11.6 | 113.7 | 69 | 136.6 | 2.2 | 141.1 |
| **105** | 89 | 65 | 90.6 | 6.1 | 102.4 | 51 | 10.8 | 0.9 | 12.7 | 65 | 90.3 | 5.1 | 100.5 | 65 | 132.0 | 2.1 | 135.4 |
| **107** | 50 | 75 | 86.4 | 5.6 | 95.3 | 61 | 11.5 | 0.8 | 13.4 | 75 | 85.4 | 4.9 | 97.1 | 75 | 139.4 | 1.3 | 141.9 |

**Supplementary Table S11. Descriptive statistics of the experimental data for the 4 analyzed traits**.

*Abbreviations: P1: parent 1, P2: parent 2, Nb: number of, SD: standard deviation, UC: usefulness criterion.*

| Trait | Year | | 2020 | | 2021 | | 2022 |
|---|---|---|---|---|---|---|---|
| | | **Site** | **CAP** | **HOU** | **CAP** | **HOU** | **HOU** |
| **Yield** | **2020** | **CAP** | | 53 | 3 | 3 | 85 |
| | | **HOU** | 0.55 | | 3 | 3 | 84 |
| | **2021** | **CAP** | 0.54 | 0.83 | | 58 | 89 |
| | | **HOU** | -0.58 | -0.20 | 0.51 | | 89 |
| | **2022** | **HOU** | 0.13 | 0.11 | 0.18 | 0.29 | |
| **Grain protein content** | **2020** | **CAP** | | 53 | 3 | 3 | 85 |
| | | **HOU** | 0.55 | | 3 | 3 | 84 |
| | **2021** | **CAP** | 0.54 | 0.83 | | 58 | 89 |
| | | **HOU** | -0.58 | -0.20 | 0.51 | | 89 |
| | **2022** | **HOU** | 0.13 | 0.11 | 0.18 | 0.29 | |
| **Plant height** | **2020** | **CAP** | | 53 | 3 | 3 | 85 |
| | | **HOU** | 0.78 | | 3 | 3 | 84 |
| | **2021** | **CAP** | 1.00 | 0.95 | | 58 | 89 |
| | | **HOU** | 0.74 | 0.56 | 0.67 | | 89 |
| | **2022** | **HOU** | 0.80 | 0.63 | 0.73 | 0.70 | |
| **Heading date** | **2020** | **CAP** | | 53 | 3 | 3 | 85 |
| | | **HOU** | 0.82 | | 3 | 3 | 84 |
| | **2021** | **CAP** | 0.78 | 0.70 | | 58 | 89 |
| | | **HOU** | 0.80 | 0.72 | 0.88 | | 89 |
| | **2022** | **HOU** | 0.79 | 0.92 | 0.92 | 0.90 | |

**Supplementary Table S12. Number of lines in common (above diagonal) and correlations (below diagonal) between sites for the 4 analyzed traits in Florimond-Desprez protocol**.

*Abbreviations for the protocol's locations are the following: CAP: Cappelle, HOU: Houville. Pearson's correlations were computed from spatially adjusted clean observations to correct the effect of the environment (location*year).*

| Trait | Year | | 2020 | | | 2021 | | | 2022 | | | |
|---|---|---|---|---|---|---|---|---|---|---|---|---|
| | | **Site** | **AUZ** | **LUS** | **MON** | **CLE** | **LUS** | **MON** | **AUZ** | **CLE** | **LUS** | **MON** |
| **Yield** | **2020** | **AUZ** | | 45 | 132 | 6 | 6 | 7 | 7 | 2 | 9 | 5 |
| | | **LUS** | 0.51 | | 132 | 41 | 41 | 35 | 40 | 16 | 34 | 37 |
| | | **MON** | 0.15 | 0.53 | | 29 | 33 | 32 | 29 | 16 | 29 | 28 |
| | **2021** | **CLE** | 0.17 | 0.37 | 0.39 | | 101 | 93 | 81 | 66 | 96 | 103 |
| | | **LUS** | 0.61 | 0.32 | 0.47 | 0.36 | | 86 | 88 | 64 | 72 | 72 |
| | | **MON** | 0.91 | 0.81 | 0.59 | 0.47 | 0.48 | | 91 | 125 | 206 | 203 |
| | **2022** | **AUZ** | 0.64 | 0.42 | 0.54 | 0.46 | 0.53 | 0.55 | | 54 | 63 | 58 |
| | | **CLE** | 1.00 | 0.55 | 0.18 | 0.33 | 0.50 | 0.28 | 0.64 | | 140 | 145 |
| | | **LUS** | 0.85 | 0.64 | 0.08 | 0.53 | 0.60 | 0.59 | 0.74 | 0.55 | | 163 |
| | | **MON** | 0.71 | 0.48 | 0.61 | 0.52 | 0.64 | 0.45 | 0.68 | 0.38 | 0.50 | |
| **Grain protein content** | **2020** | **AUZ** | | 45 | 128 | 7 | 6 | 7 | | | | |
| | | **LUS** | 0.58 | | 131 | 41 | 40 | 35 | | | | |
| | | **MON** | 0.59 | 0.70 | | 29 | 32 | 32 | | | | |
| | **2021** | **CLE** | 0.62 | 0.54 | 0.53 | | 100 | 93 | | | | |
| | | **LUS** | 0.26 | 0.62 | 0.61 | 0.75 | | 86 | | | | |
| | | **MON** | 0.06 | 0.80 | 0.63 | 0.39 | 0.51 | | | | | |
| **Plant height** | **2020** | **AUZ** | | 46 | 132 | 7 | 6 | 7 | 7 | 2 | 10 | 5 |
| | | **LUS** | 0.91 | | 132 | 41 | 41 | 35 | 40 | 16 | 34 | 37 |
| | | **MON** | 0.84 | 0.85 | | 29 | 33 | 32 | 29 | 16 | 29 | 28 |
| | **2021** | **CLE** | 0.93 | 0.79 | 0.80 | | 102 | 94 | 83 | 66 | 96 | 105 |
| | | **LUS** | 0.76 | 0.84 | 0.87 | 0.81 | | 87 | 88 | 64 | 72 | 72 |
| | | **MON** | 0.94 | 0.56 | 0.79 | 0.76 | 0.83 | | 91 | 125 | 208 | 204 |
| | **2022** | **AUZ** | 0.98 | 0.93 | 0.92 | 0.53 | 0.77 | 0.83 | | 54 | 63 | 58 |
| | | **CLE** | 1.00 | 0.86 | 0.74 | 0.27 | 0.52 | 0.58 | 0.63 | | 140 | 146 |
| | | **LUS** | 0.94 | 0.86 | 0.69 | 0.66 | 0.82 | 0.82 | 0.76 | 0.71 | | 165 |
| | | **MON** | 0.98 | 0.90 | 0.91 | 0.69 | 0.85 | 0.84 | 0.78 | 0.80 | 0.83 | |
| **Heading date** | **2020** | **AUZ** | | 46 | 132 | 7 | 6 | 7 | 7 | 2 | 10 | 5 |
| | | **LUS** | 0.91 | | 131 | 41 | 41 | 34 | 40 | 16 | 34 | 36 |
| | | **MON** | 0.53 | 0.86 | | 29 | 33 | 32 | 29 | 16 | 29 | 28 |
| | **2021** | **CLE** | 0.90 | 0.88 | 0.79 | | 102 | 94 | 83 | 66 | 96 | 105 |
| | | **LUS** | 0.90 | 0.91 | 0.88 | 0.91 | | 87 | 88 | 64 | 72 | 72 |
| | | **MON** | 0.85 | 0.96 | 0.91 | 0.91 | 0.95 | | 91 | 125 | 208 | 204 |
| | **2022** | **AUZ** | 0.99 | 0.94 | 0.85 | 0.88 | 0.94 | 0.90 | | 54 | 63 | 58 |
| | | **CLE** | 1.00 | 0.88 | 0.67 | 0.92 | 0.92 | 0.92 | 0.91 | | 140 | 146 |
| | | **LUS** | 0.83 | 0.95 | 0.87 | 0.84 | 0.90 | 0.92 | 0.91 | 0.94 | | 165 |
| | | **MON** | 0.95 | 0.94 | 0.92 | 0.91 | 0.93 | 0.94 | 0.94 | 0.96 | 0.96 | |

**Supplementary Table S13. Number of lines in common (above diagonal) and correlations (below diagonal) between sites for the 4 analyzed traits in INRAE protocol**. *Abbreviations for the protocol's locations are the following: AUZ: Auzeville, CLE: Clermont-Ferrand, LUS: Lusignan, MON: Mons. Pearson's correlations were computed from spatially adjusted clean observations to correct the effect of the environment (location*year).*
